# Vps35 p.D620N causes Lrrk2 kinase hyperactivity, chronic microglial activation and inflammation

**DOI:** 10.64898/2026.03.09.710482

**Authors:** Isaac Bul Deng, Mengfei Bu, Jordan Follett, Robert C. Sharp, Adamantios Mamais, Leyna Xoi, Fahong Yu, Georges Rabil, Shannon Wall, Matthew J. Farrer

**Affiliations:** Department of Neurology, McKnight Brain Institute, University of Florida, Gainesville, Florida, USA; Viral Immunology and Intravital Imaging Section, National Institute of Neurological Disorders and Stroke (NINDS), National Institutes of Health (NIH), Bethesda, Maryland, USA; Interdisciplinary Center for Biotechnology Research, University of Florida, Gainesville, Florida, USA; Department of Biostatistics, University of Florida, Gainesville, Florida, USA

**Author notes:** To whom the correspondence should be addressed: Isaac Bul Deng and Matthew J. Farrer. Contributed equally to the study (Isaac Bul Deng, Mengfei Bu).

## Abstract

Pathogenic variants in *leucine-rich repeat kinase 2* (*LRRK2*), *RAB32*, and *vacuolar protein sorting 35* (*VPS35*) cause dominantly inherited, late-onset Parkinson’s disease (PD). Mutations in all three genes constitutively activate LRRK2 kinase activity, enhancing innate immune defense against pathogens; however, chronic activation is linked to dopaminergic neuronal degeneration. LRRK2 and RAB32 are highly expressed in all myeloid cells including brain microglia, whereas VPS35, a core component of the retromer, is ubiquitously expressed. Nevertheless, the VPS35 p.D620N substitution induces the greatest constitutive increase in endogenous LRRK2 kinase activity, and its consequences for immune function remain unclear. We therefore examined the transcriptomic and functional effects of wild type (WT) and Vps35 p.D620N knock-in mice (VKI) from microglia, isolated from brains of naïve animals at six-month of age, and initially analyzed by single-cell RNA sequencing (scRNAseq). Despite the absence of overt behavioral differences, differential gene expression revealed genotype-dependent changes in antimicrobial humoral immunity, lysosomal stress sensing, and phagocytosis. *S100 family genes* and *lipocalin 2* were markedly upregulated, a finding subsequently validated by gene expression and immunohistochemistry. Downregulation was also observed in pathways governing microglial regulation of synaptic transmission, neuronal development and homeostatic immune signaling. Peripherally-administered lipopolysaccharide (LPS) stimulated microglial activation and phagocytic markers in WT brain, and further increased morphological activation and synaptic engulfment in VKI animals. Compared to WT, Vps35 p.D620N drives a chronic pro-inflammatory microglial state characterized by elevated innate immune signaling, lysosomal stress and enhanced phagocytic activity. Hence, heightened microglial sensitivity, further exacerbated by a peripheral immune challenge, offers a parsimonious, if multicellular, explanation for the multifactorial etiology of PD, and may help to account for its penetrance and variable expressivity. Together, these findings provide mechanistic insight into how retromer dysfunction and LRRK2 kinase hyperactivity intersect with microglial biology to influence PD pathogenesis, potentially promoting synaptic remodeling and neurodegenerative vulnerability.

## INTRODUCTION

Parkinson’s disease (PD) is defined by motor dysfunction resulting from dopamine deficiency associated with degeneration of midbrain dopaminergic (DA) neurons in the *substantia nigra pars compacta* (*SNpc*).^1,2^ It is the most common neurodegenerative movement disorder, affecting approximately twelve-million people world-wide. Patients suffer a constellation of non-motor symptoms, many of which are unresponsive to dopaminergic replacement therapy and are equally debilitating as the preceding onset of the motor symptoms. While much of the etiology of PD remains elusive, it is considered a complex interaction between aging, environmental factors and genetic predisposition.^3^ Monogenic factors now account for 5-10% of cases, while the remainder have no known cause (idiopathic). Of known genetic causes, mutations in *leucine-rich repeat kinase 2* (*LRRK2*) confer the greatest genotypic and population-attributable risk.^4^ The most frequent of these variants, LRRK2 Gly2019Ser and its geographical enrichment is driven by positive selection^5^, as it promotes proinflammatory immune responses in cells of myeloid-lineage.^6,7^ However, those responses lead to an increase in age-associated PD, a concept known as antagonistic pleiotropy.^4^

*SNpc* neurons are selectively vulnerable given their specialized morphology, post-mitotic microenvironment and melanocyte biology required to produce, package, release and reuptake dopamine. Maintaining a healthy proteome with such complex morphology distal to the cell soma is likely to predispose DA neurons to lysosomal and mitochondrial stress, and homeostasis requires both cell-autonomous and non-cell autonomous factors.^1,2^ *Post-mortem* gliosis in the *SNpc* has long been observed, as both astroglia and microglia are reactive, and microglia are regionally enriched relative to other brain nuclei.^2,3,8^ Microglia continuously survey the parenchyma with their motile projections and transition into states that facilitate clearance of tau and alpha-synuclein aggregates associated with PD and other common neurodegenerative diseases.^9–12^ Unlike astrocytes and oligodendrocytes, which can be replenished through gliogenesis in adulthood from radial glial cells, microglia originate from myeloid precursors in the yolk sac, not the neuroectoderm. Microglia progenitor cells migrate into the brain parenchyma during development, before the formation of the blood–brain barrier. Once settled in the brain, they mature and persist throughout an individual’s lifetime. Although microglia can be replenished through self-renewal, their capacity for repopulation is limited.^13,14^ Recent advances in transcriptomic analysis of central nervous system immune cells has uncovered microglial subsets including disease-associated microglia (DAMs), primarily deemed to contain and remove damage, with capabilities of either ameliorating or exacerbating disease progression, with other subtypes implicated in stress, cytokine/ chemokine signalling^15–17^ or senescence.^18^ Our group discovered pathogenic variants in *leucine-rich repeat kinase 2 (LRRK2),*^19–22^ *RAB32,* ^23^ and *vacuolar protein sorting 35 (VPS35*)^24,25^ in families with dominantly-inherited parkinsonism, in which disease is clinically (and often pathologically) indistinguishable from idiopathic late-onset PD. All three causes activate LRRK2 kinase, and these proteins appear to physically interact, along with RAB29, in endolysosomal sorting, recycling and lysosome-related cargo degradation.^26,27^

We and others have characterized Vps35 p.D620N knock-in mice (VKI) to discover they possess the highest constitutive increase in endogenous Lrrk2 kinase activity, even when compared to pathogenic mutant models of *Lrrk2* or *Rab32.*^23,28–30^ VKI mice also exhibit a significant reduction in the density of striatal dopamine transporter (Dat), and increased evoked striatal dopamine release, all of which are restored via inhibition of Lrrk2 kinase activity with MLi-2 for 7-days.^30^ These findings indicate a functional converge for Vps35 and Lrrk2 that warrants further investigation. In WT mice, we discovered that peripheral immune stimulation acutely drives Rab32 expression in microglia that correlates with Lrrk2 kinase activity.^31^ Epidemiologic and modeling data also suggests LRRK2 disease penetrance is a function of immune exposure.^32,33^ Congruent with these findings, LRRK2 kinase activity may be elevated in idiopathic PD.^31^ Hence, LRRK2 kinase inhibitors are in clinical trials as disease-modifying therapy for PD. More mechanistic insight is evidently needed on the role of microglial Lrrk2-Vps35 in inflammation, which can inform microglia targeted gene therapy for PD. Microglia in the central nervous system (CNS) specifically express triggering receptor expressed on myeloid cells (TREM2), which is recycled back to the plasma membrane via the retromer (VPS35) complex.^32^ TREM2 has anti-inflammatory properties, and its genetic variants Arg47His and Arg62His have been linked to Alzheimer’s disease (AD).^33^ Lrrk2 kinase inhibition restores the levels of Vps35 cargo, Dat^30^, and extending this knowledge to Trem2 may yield important insights for PD and AD. In this study we explored the transcriptomic profile of enriched microglia obtained from Vps35 p.D620N mice, and its relationship with microglial activation, phagocytic capacity and synaptic pruning. Our data show that Vps35 p.D620N drives a chronic pro-inflammatory microglial state characterized by elevated innate immune signaling, lysosomal stress, and enhanced phagocytic activity.

## RESULTS

### Single cell RNA sequencing (scRNAseq) on enriched brain microglia

To elucidate the effects of Vps35 p.D620N on the microglia-specific transcriptome, we analyzed purified CD11b+ myeloid cells obtained from 6-month-old VKI mice and WT littermates using 10x Genomics ScRNAseq. A total of 120,730 cells were analyzed after quality control. Clustrees analysis at 0.5 resolution differentiated the major cell types (Fig. 1a), defined using the CellMarker2.0 database. Differential counts within clusters were not significantly different by genotype. Canonical sets of genes for each cell type were collated based on mouse scRNAseq literature^34–36^, and their expression was confirmed to verify CellMarker database cluster prediction (Supplemental Fig. 1a). A Umap plot of different cell types from WT, VKI^Het^ and VKI^Hom^ was generated with Seurat at 0.5 resolution, revealing 24 distinct clusters (Fig. 1b); 68.58% of the cells analyzed were microglia with others prevalent as described in Fig. 1c. Using pair-wise analysis (adjusted p-value < 0.05 and fold change > 1.4), a total of 96 genes were upregulated, and 538 downregulated in VKI^Het^ compared to WT. Similarly, 121 genes were upregulated, and 502 downregulated in VKI^Hom^ compared to WT. Gene ontology revealed enrichment of pathways involved in acute inflammation, complement activation, lysosomal stress/phagocytosis, tissue remodeling, lipid metabolism and cell proliferation in VKI^Het^ versus WT (Supplemental Fig. 1b, c, Supplemental Fig. 2). These pathways were further enriched in VKI^Hom^ with enhanced inflammatory phenotypes (Fig. 1d, e, Supplemental Fig. 3). Moreover, there was downregulation in pathways enriched for synaptic transmission, synaptic development/ modelling and gliogenesis in VKI^Het^(Supplemental Fig. 1b, d, Supplemental Fig. 4), with greater neuronal/glial suppression observed in VKI^Hom^ (Fig. 1f, Supplemental Fig. 5). Clusters 0, 1 & 2 in the Umap (Fig. 1b) were identified as microglia, our primary focus; thus, we regrouped them bioinformatically removing other cell types. Clustrees analysis (Fig. 2a) at 0.5 resolution yielded 6 unique microglia subclusters, henceforth denoted by MG and followed by cluster number (Fig. 2b). The majority of the cells expressed classical homeostatic microglial markers, such as *Cx3cr1*, *Tmem119*, *Hexb*, *P2ry12*, *P2ry13* and *Sall1* (Fig. 2c). There were no substantial differences in the cell count in the clusters by genotype (Fig. 2d, e). MG0 and 4 were identified as quiescent microglia, as indicated by their rich expression of homeostatic microglial genes, with concomitant downregulation of immune/cell proliferative and ribosomal genes (Fig. 2e, Supplemental Fig. 6). Additionally, MG1 and 3 were identified as interferon-responsive microglia (*Stat1, Stat2*, *Arhgap15, Ifitm3*), and MG2 as translationally primed/activated microglia exemplified by the prominent expression of ribosomal genes (e.g. *Rps26*). The transcriptomic signature of MG5 resembles intermediary disease-associated microglia (DAMs), with defining genes such as *Apoe, Cd68, Ctsd, S100a8 and S100a9,* although they have yet to fully transform as denoted by their partial expression of homeostatic microglial genes *(Gpr34, P2ry13, Hexb, Slc2a5)* (Fig. 2e, Supplemental Fig. 6).

**Fig. 1.**
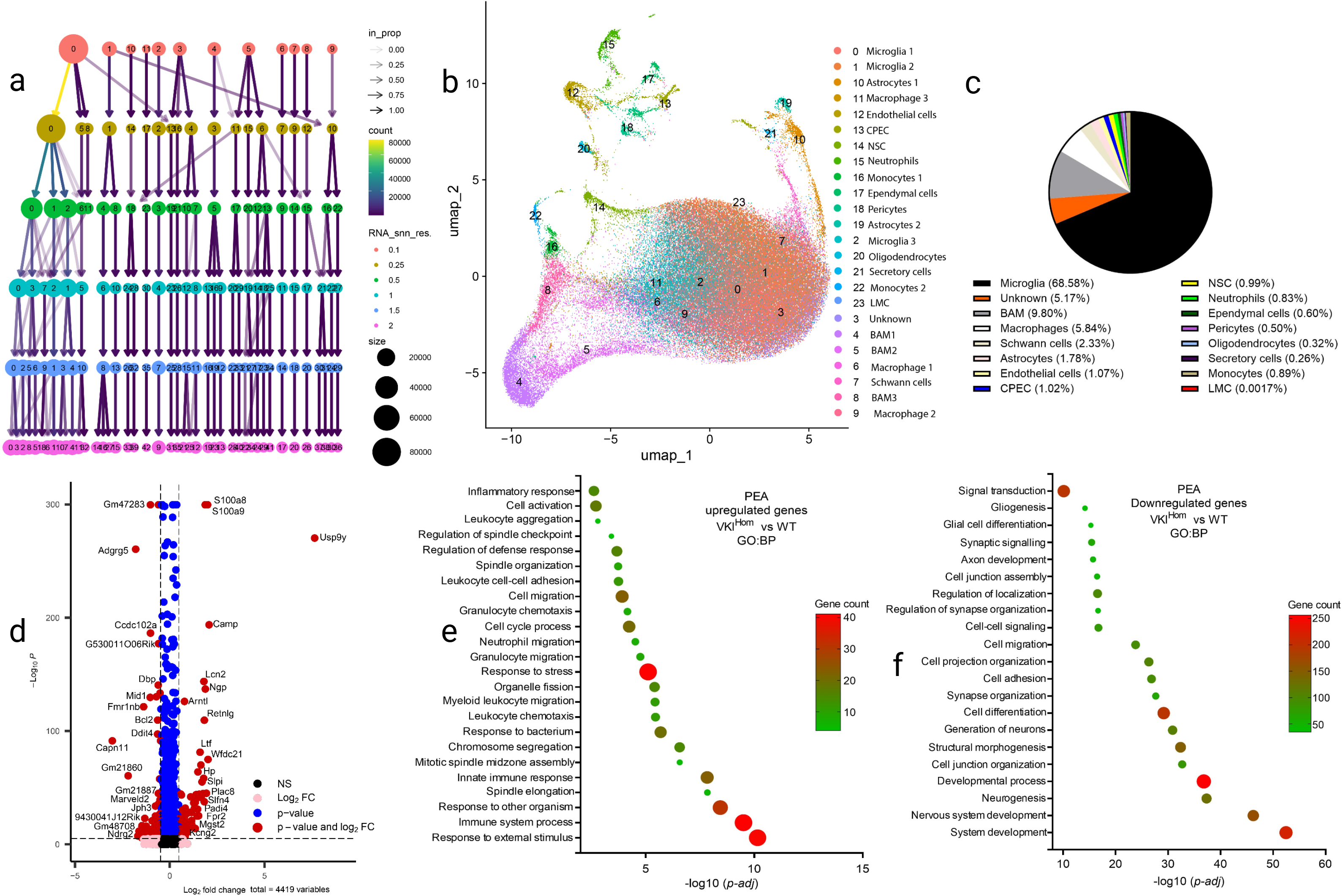
10x Genomics scRNAseq of CD11b^+^ antibody-based captured microglia from whole brain of VKI mice at ∼6M. **a**) The clustrees analysis at 0.5 resolution differentiated most major cell types, defined using CellMarkers, although differential cell counts within clusters were not significantly different by genotype (not shown). **b**) Umap plot of different cell types following CD11b^+^ antibody-based selection (WT n = 2, VKI^Het^ n = 3 and VKI^Hom^ n = 3) (120,730 cells) at 0.5 resolution. **c**) Proportion of cell types analyzed, >68% microglia. **d**) Volcano plot generated with the EnhancedVolcano R package for upregulated and downregulated genes for all cell types in VKI^Hom^. **e, f**) g:Profiler g:GOSt pathway enrichment analysis for upregulated and downregulated genes for all the cell types in VKI^Hom^. Abbreviations: border-associated macrophages (BAM), choroid plexus epithelial cells (CPEC), neural stem cells (NSC) and leptomeningeal cells (LMC).

**Fig. 2.**
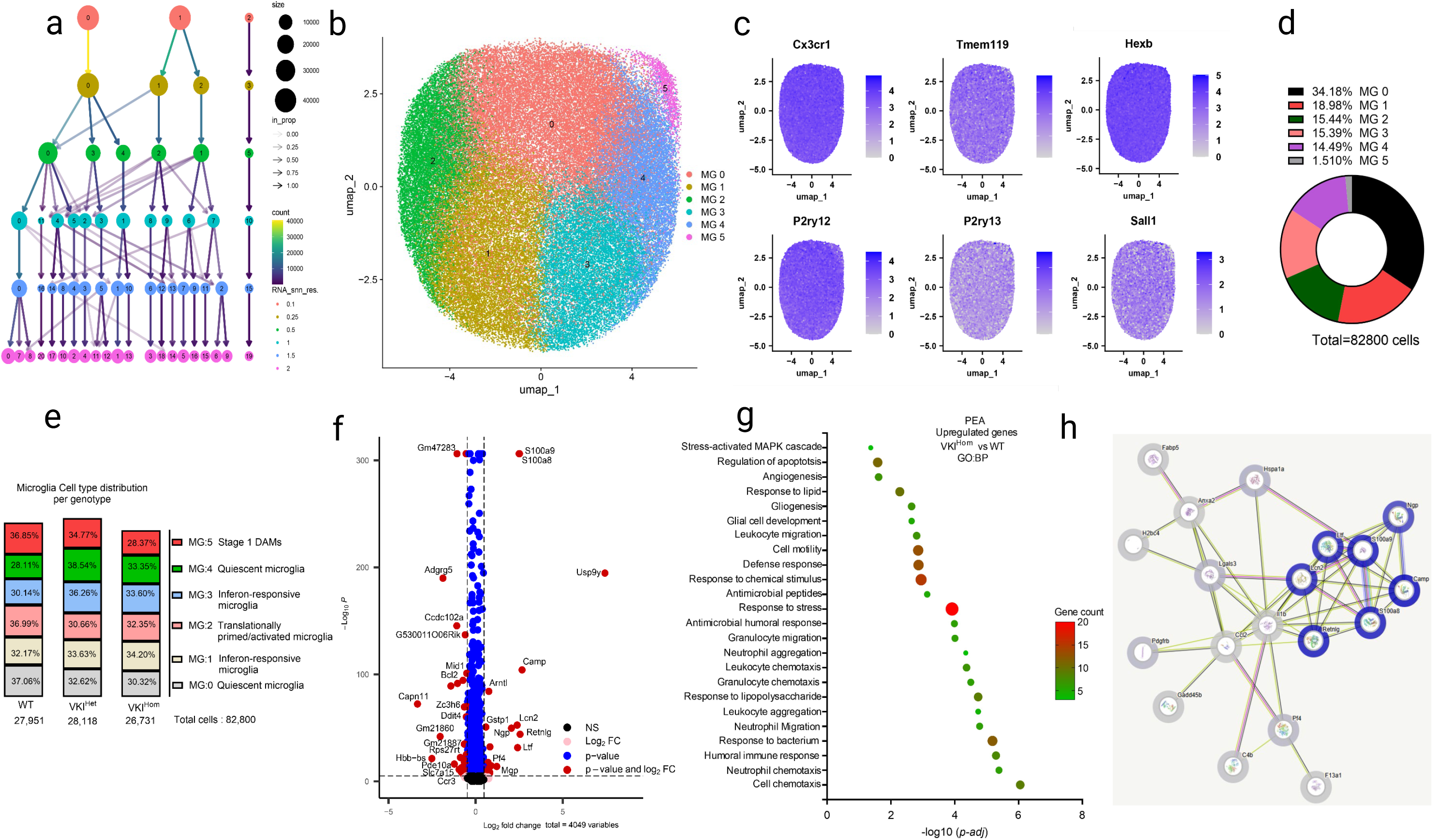
Vps35 p.D620N increases expression of microglial genes involved in innate immunity/acute inflammation, phagocytosis and lysosomal stress. **a**) Clustrees re-analysis of microglia clusters 0, 1 & 2 from Fig. 1b at 0.5 resolution. **b**) Umap corresponding to these microglia subclusters (total cells: 82,800). **c**) Expression of classical microglial gene markers in the subclusters. **d**) Cell count distribution for the subclusters. **e**) Microglia cell type distribution per genotype per a subcluster. **f**) Volcano plot generated with the EnhancedVolcano R package for upregulated, and downregulated genes in VKI^Hom^ microglia. **g**) g:Profiler g:GOSt pathway enrichment analysis for upregulated genes in VKI^Hom^ microglia. **h**) STRING generated protein-protein interaction network and functional enrichment analysis for upregulated genes in VKI^Hom^, with disconnected nodes excluded from network display.

### Vps35 p.D620N increases expression of microglial genes involved in innate immunity/acute inflammation, phagocytosis and lysosomal stress

Next, we used Seurat for differential gene expression analysis of microglia cells and generated volcano plots (Fig. 2f). Pair-wise analysis (adjusted p-value < 0.05 and fold change > 1.4) revealed a total of 36 upregulated and 39 downregulated genes in VKI^Het^ compared to WT. In contrast, there were 49 upregulated and 78 downregulated genes in VKI^Hom^ compared to WT. Based on STRING analysis supplemented with g:Profiler g:GOSt pathway enrichment analysis (PEA), VKI^Hom^ microglia compared to WT had genes enriched for acute inflammation, nuclear factor kappa B (NFκB) and interleukin 1β pathways. The latter was noted by marked expression of chemokine and antimicrobial/alarmin genes including *Il1b, Ccl2, Lcn2, S100a8, S100a9, Retnlg, Camp, Ltf and Ngp* (Fig. 2f - h), a feature of microglial activation. This phenotype was also observable in VKI^Het^ (Supplemental Fig. 7b, c, d). Moreover, in VKI^Hom^, there was enrichment for genes involved in lysosomal stress/damaging sensing and phagocytosis (*Lgals3, Anxa2*), complement/innate effector engagement (*C4b, Pf4*), cellular stress/heat shock response (*Hspa1a and Gadd45b*) and interferon restriction factor with immune transcriptional modulators and a circadian regulator (*Ifitm2, Nfil3 and Arntl*) (Fig. 2f - g). These findings suggest endolysosomal dysfunction/damaging sensing in microglia, consistent with retromer (Vps35) impairment, and complement engagement which coincides with immune activation, clearance, and neuroinflammatory amplification, as well as proteostatic/metabolic stress and immune-programing,accompanied inflammation. Compared to WT, VKI^Hom^ had downregulation in G-protein coupled receptors and chemokine receptors (*Adgrg5, Ccr3, Ccl5 and Gpr68*) involved in environmental sensing and immune tuning, suggesting suppression of microglial homeostatic signaling capacity (Supplemental Fig. 7a). The latter changes were accompanied by downregulation of metabolic/mitochondrial bioenergetic support *(Slc25a54, Abca13, Pde10a, Rps27rt*), and related cytoskeletal transport with impaired skeleton/scaffold regulation (*Tubb1, Rack1, Mid1, Frmp1b*) (Supplemental Fig. 7a). Similarly, in VKI^Het^, there was downregulation in genes implicated in immune modulation and remodeling of the cytoskeleton and synapses (Supplemental Fig. 7b, e, f).

### Validation of scRNAseq humoral immune antimicrobial peptides

The most striking revelation of scRNAseq was the upregulation of humoral antimicrobial peptides in naïve VKI microglia, prominently the S100 calcium binding protein family (S100), and lipocalin 2 (Lcn2). Thus, we used immunofluorescence (IHC), and real-time quantitative polymerase chain reaction (RT-qPCR) to validate the expression of *S100a9* and *Lcn2*, both of which are potent antimicrobial peptides with strong connection to injury and neurodegeneration.^37–40^ These experiments were carried out using frozen tissues from midbrain as it is the most vulnerable/relevant region to idiopathic PD. IHC with confocal microscopy confirmed the significant upregulation of microglial S100a9 (Fig. 3a, b) and Lcn2 (Fig. 3d, e) in naïve VKI mice. However, there was only a modest difference using RT-qPCR for *S100a9* (Fig. 3c) and *Lcn2* (Fig. 3f), although these divergent results may be attributable to the small sample size. Antimicrobial peptides are highly secreted in response to microbial challenges. Therefore, as a proof of concept, we injected naïve WT mice with a daily intraperitoneal dose of 1 mg/kg of LPS for 2 days. We have shown this LPS dosage upregulates midbrain expression of microglial Rab32, with concomitant activation of Lrrk2 kinase activity, which we consider essential in microgliosis.^41^ Using this paradigm, IHC confirmed that LPS potently upregulates microglial S100a9 (Fig. 3g, h) and Lcn2 (Fig. 3j, k), with corresponding increases in their mRNA levels in the midbrain tissue, respectively (Fig. 3i, l).^40,42^

**Fig. 3.**
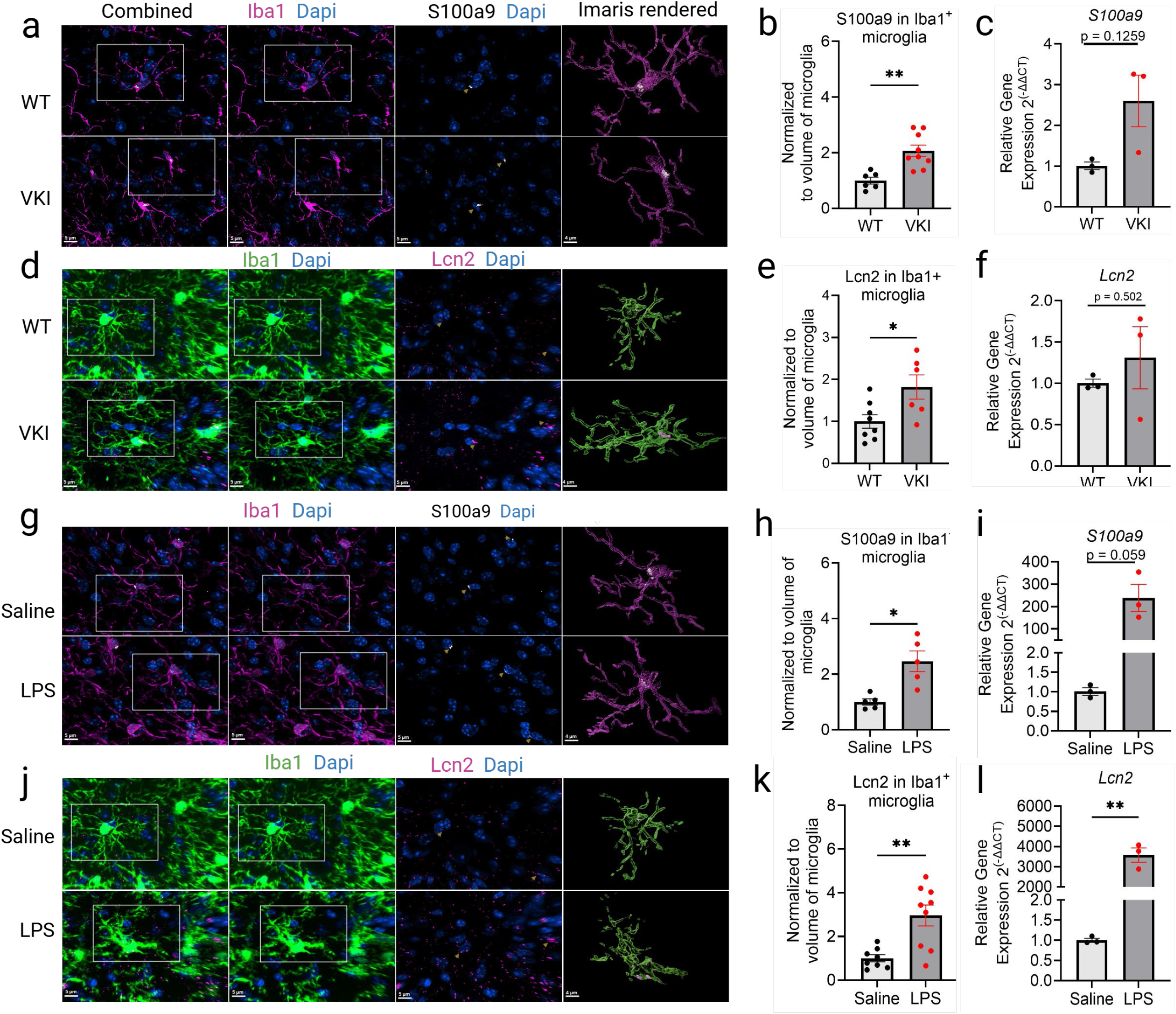
Vps35 p.D620N increases microglial expression of humoral immune antimicrobial peptides, S100a9 and Lcn2. a, d, g, **j**) Immunofluorescence confocal microscopy (IF-CM) representative images for S100a9 and Lcn2 co-stained with Dapi, and their Imaris^TM^ rendering in the midbrain of naïve VKI^Hom^, and in WT mice injected with LPS. **b**) Volumetric analysis of S100a9^+^ puncta in Iba1+ microglia in WT vs VKI^Hom^ (unpaired *t*-test; **p = 0.0017). [WT n = 4(6); VKI^Hom^ = 5(9)]. **c**) RT-qPCR relative gene expression of *S100a9* in WT versus VKI^Hom^ (unpaired *t*-test, p = 0.1259; n = 3 per a group). **e**) Volumetric analyses of Lcn2^+^ puncta in Iba1+ microglia in WT vs VKI^Hom^ (unpaired *t*-test, *p = 0.0384; WT n = 4 (8); VKI^Hom^ = 3 (6). **f**) RT-qPCR relative gene expression analysis of *Lcn2* in WT vs VKI^Hom^ (unpaired *t*-test; p = 0.5019; n = 3 per a group). **h**) Volumetric analysis of S100a9^+^ puncta in Iba1+ microglia in Saline versus LPS injected WT mice (unpaired *t*-test with Welch’s correction, *p = 0.0142). [Saline n = 4(5); LPS = 5]. **i**) RT-qPCR relative gene expression of *S100a9* in saline versus LPS injected WT mice (unpaired *t*-test, p = 0.0591; n = 3 per a group). **k**) Volumetric analysis of Lcn2^+^ puncta in Iba1^+^ microglia in saline versus LPS injected WT mice (unpaired *t*-test with Welch’s correction, **p = 0.0031). [Saline n = 4 (8); LPS = 5 (9)]. **l**) RT-qPCR relative gene expression of *Lcn2* in saline versus LPS injected WT mice (unpaired *t*-test, **p = 0.0098; n = 3 per group). *p <0.05, **p<0.01. Error bars are ± SEM. n refers to the number of mice, and in parentheses the number of tissue sections. The yellow arrow denotes the punctas.

### Vps35 p.D620N exacerbates inflammation and phagocytosis under inflammatory conditions in the midbrain

The scRNAseq in VKI microglia revealed elevated expression in genes involved in innate immunity/acute inflammation, phagocytosis and lysosomal stress. As such, we postulated that acute inflammation may exacerbate these states in a manner that will trigger early onset of cardinal pathological features of PD such as loss of midbrain DA neurons^43,44^ and will be most pronounced in VKI mice relative to WT littermates. In VKI mice, these neurodegenerative pathologies are noted by 14 - 16 months of age, albeit inconsistently.^43–46^ Congruent with our recent studies, we confirmed that Vps35 p.D620N doesn’t alter the expression of Vps35 protein, and other members of the retromer complex such as Vps26 (Fig. 4a, f, g). VKI mice have no overt loss of DA neurons in the MB at 6M as exemplified by Th protein expression, albeit a gross measure. Systemic inflammation in VKI mice evoked a modest increase in Iba1; nonetheless, the inflammation appears augmented in these mice as shown by the protein expression of Stat1 and Cd11b (Fig. 4 a, c–e). By IHC, microglial Trem2 expression was strongly suppressed by LPS-induced inflammation with effects more pronounced in VKI mice (Fig. 4b, h, j). These results coincided with microglial expression of Cd68, a marker of their activation and phagocytosis. As a control for retromer function, we examined similar markers in Vps35 haploinsufficient mice (50% reduction in Vps35) (cVKI). Reduction of Vps35 protein (Fig. 5a, f) destabilizes the retromer complex as shown by the concomitant reduction of Vps26 (Fig. 5a, g). Under inflammatory conditions, cVKI mice had a greater Trem2 suppression (Fig. 5b), accompanied by exacerbated neuroinflammation as depicted by the protein levels of Iba1, Stat1 and Cd11b. However, this expression profile did not directly correlate with IHC of microglial expression of Trem2 and Cd68 (Fig. 5h–k). In addition, LPS significantly elevated the expression of lipoprotein lipase (LPL) in midbrain, a protein with critical roles in lipid metabolism, and a marker of activated/phagocytic microglia, without genotypic differences (Fig.8a –d ).

**Fig. 4.**
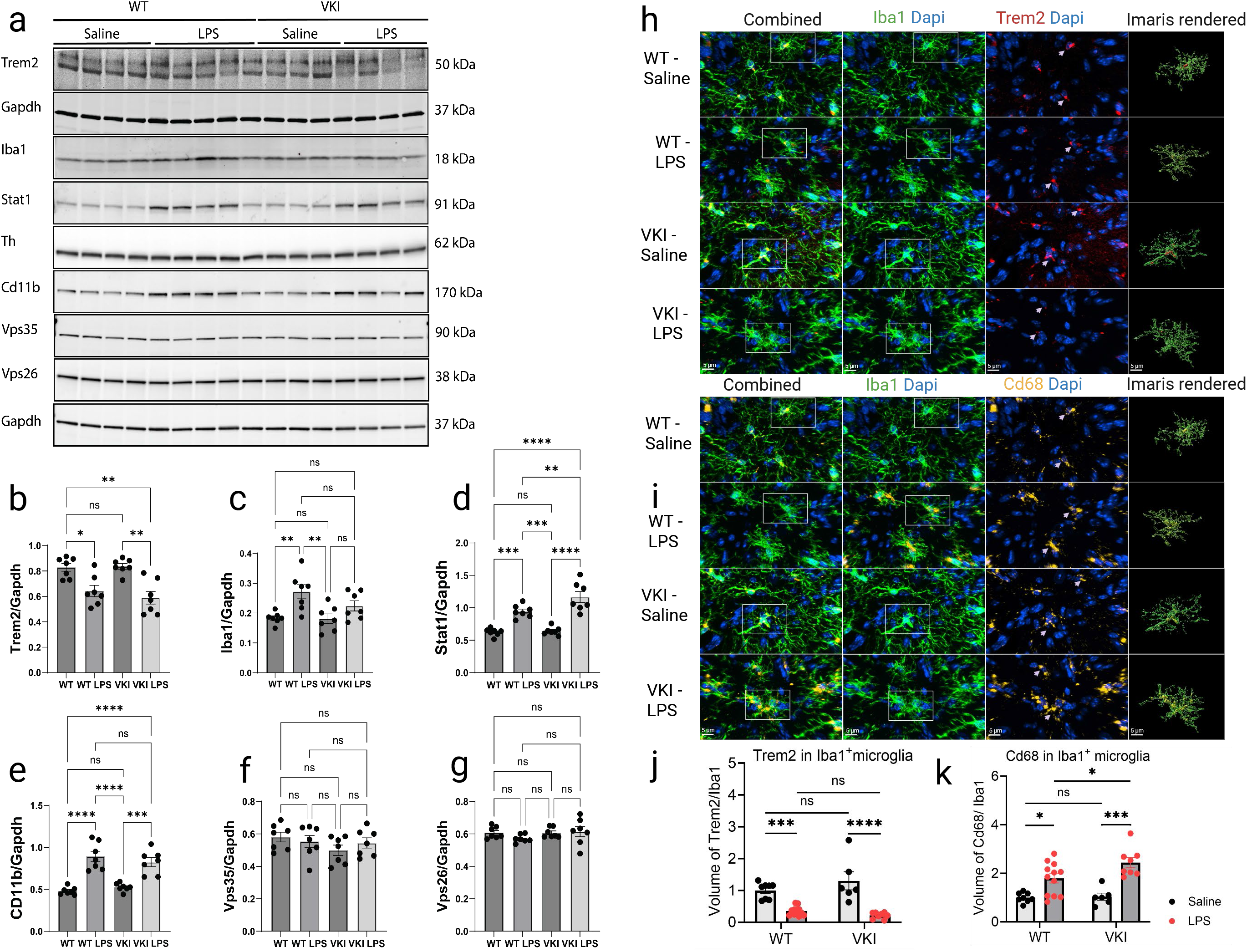
Vps35 p.D620N exacerbates inflammation and phagocytosis under inflammatory conditions in the midbrain. **a)** Representative immunoblots for Trem2, Iba1, Stat1, Th, Cd11b, Vps35 and Vps26 in the midbrain. Densitometric analyses by two-way ANOVA with Tukey’s multiple comparisons test of: Trem2 **(b**, WT vs WT LPS, *p = 0.0149; WT vs VKI LPS, **p = 0.0018; VKI vs VKI LPS, **p = 0.0013), Iba1 **(c**, WT vs WT LPS, **p = 0.0071; VKI vs WT LPS, **p = 0.0060), Stat1**(d,** WT vs WT LPS, ***p = 0.0001; WT vs VKI LPS, ***p <0.0001; WT LPS vs VKI, ***p = 0.0003; VKI vs VKI LPS, ****p <0.0001; WT LPS vs VKI LPS, **p = 0.0056), Cd11b **(e**, WT vs WT LPS, ****p<0.0001; WT vs VKI LPS, ****p <0.0001; VKI vs WT LPS, ****p <0.0001; VKI vs VKI LPS, ***p = 0.0002), Vps35 **(f)** and Vps26 **(g)**. n = 7 for each group. **h, i**) IF-CM for Iba1, Trem2 and Cd68 co-stained with Dapi, and their Imaris^TM^ rendering in the midbrain of WT and VKI^Hom^ mice, with/without LPS. Two-way ANOVA with Tukey’s multiple comparisons test of: Trem2^+^ puncta in Iba1^+^ microglia (**j**, WT vs WT LPS, ***p = 0.0008;VKI vs VKI LPS, ****p<0.0001) and Cd68^+^ puncta in Iba1+ microglia (**k**, WT vs WT LPS, *p = 0.0120; WT LPS vs VKI LPS, *p = 0.0475; VKI vs VKI LPS, ***p = 0.0001). *p <0.05, **p<0.01, *\*\*\*p*<0.001, \**\*\*\*p*<0.0001. Error bars are ± SEM. [WT n = 4 (8); WT LPS n = 6 (12); VKI^Hom^ Saline n = 3(6); VKI^Hom^ LPS = 4(8)]. n refers to the number of mice and in parentheses the number of tissue sections.

**Fig. 5.**
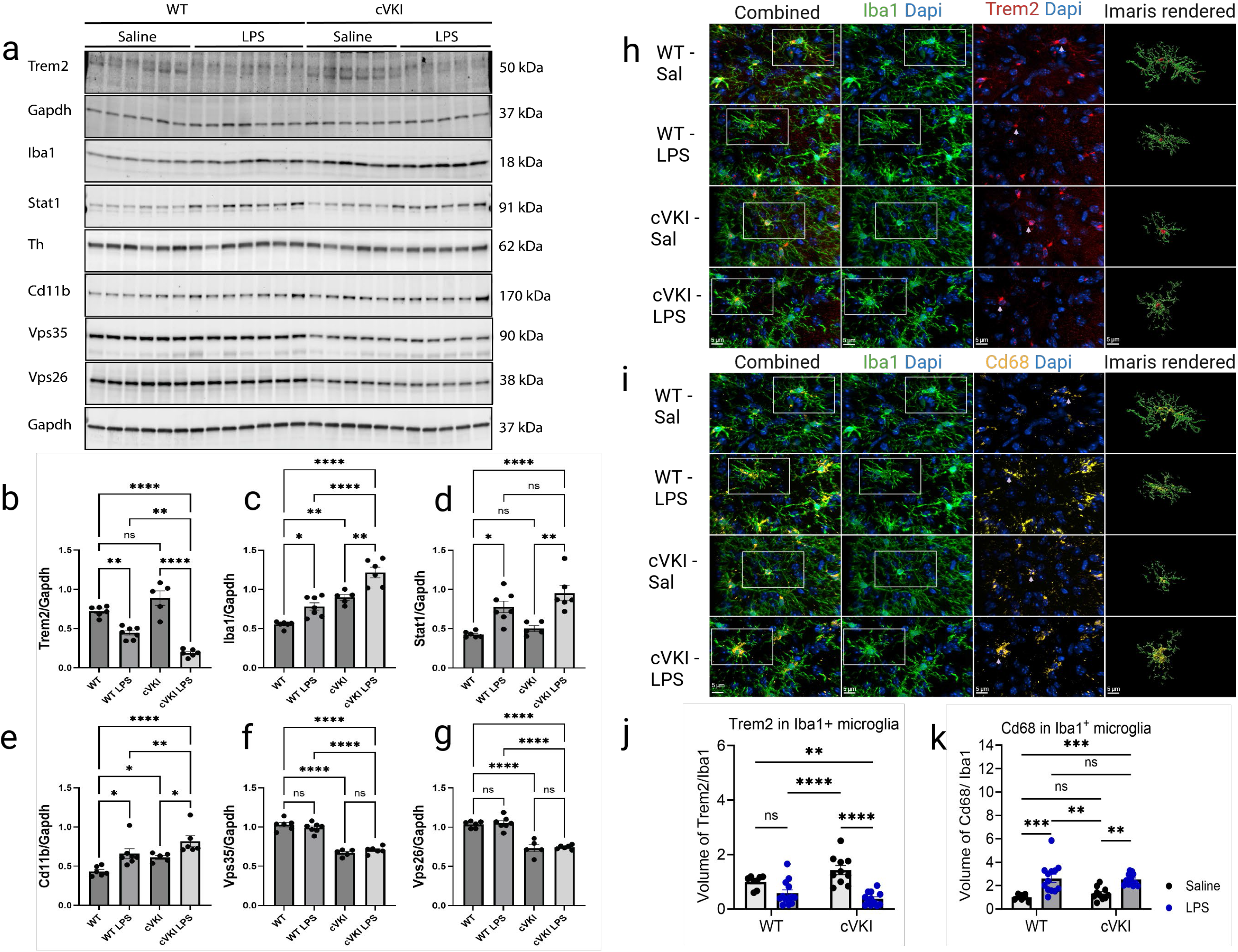
Vps35 haploinsufficiency potentiates microglial activation in the midbrain under inflammatory conditions. **a)** Representative immunoblots for Trem2, Iba1, Stat1, Th, Cd11b, Vps35 and Vps26 in the midbrain. Densitometric analyses by two-way ANOVA with Tukey’s multiple comparisons test of: Trem2 **(b**, WT vs WT LPS, **p = 0.0059; WT vs cVKI LPS, ****p = <0.0001; cVKI vs cVKI LPS ****p<0.0001; WT LPS vs cVKI LPS **p = 0.0086), Iba1 **(c**, WT vs WT LPS, *p = 0.0430; WT vs cVKI, **p = 0.0012; WT vs cVKI LPS, ****p<0.0001; WT LPS vs cVKI LPS, ****p <0.0001; cVKI vs cVKI LPS, **p = 0.0046), Stat1**(d,** WT vs WT LPS, *p = 0.0126; WT vs cVKI LPS, ****p<0.0001, cVKI vs cVKI LPS, **p = 0.0013), Cd11b **(e**, WT vs WT LPS, *p = 0.0346; WT vs cVKI LPS, ****p<0.0001; WT LPS vs cVKI LPS, **p = 0.0053; WT vs cVKI, *p = 0.0129; cVKI vs cVKI LPS, *p = 0.0324), Vps35 **(f,** WT vs cVKI, ****p<0.0001; WT vs cVKI LPS, ****p<0.0001; WT LPS vs cVKI LPS,****p<0.0001) and Vps26 **(g,** WT vs cVKI, ****p<0.0001; WT vs cVKI LPS, ****p<0.0001; WT LPS vs cVKI LPS,****p<0.0001). WT n = 6; WT LPS n = 7; cVKI n = 5; cVKI LPS =6. **h, i**) IF-CM for Iba1, Trem2 and Cd68 co-stained with Dapi, and their Imaris^TM^ rendering in the midbrain of WT and cVKI mice, with/without LPS. Volumetric analyses by two-way ANOVA with Tukey’s multiple comparisons test of: Trem2^+^ puncta in Iba1+ microglia (**j**, WT vs cVKI LPS, **p = 0.0067; WT LPS vs cVKI ****p<0.0001; cVKI vs cVKI LPS,****p<0.0001) and Cd68^+^ puncta in Iba1+ microglia (**k**, WT vs WT LPS, ***p = 0.0003; WT vs cVKI LPS,***p = 0.0010; WT LPS vs cVKI, **p = 0.0021;cVKI vs cVKI LPS, **p = 0.0060). *p <0.05, **p<0.01, *\*\*\*p*<0.001, \**\*\*\*p*<0.0001. Error bars are ± SEM. [WT n = 5 (10); WT LPS n = 7(14); cVKI n = 5(10); VKI LPS = 6(12)]. n refers to the number of mice and in parentheses the number of tissue sections.

### Vps35 p.D620N exacerbates microglial activation and phagocytosis under inflammatory conditions in striatum, and may potentiate their pruning of synapses

Midbrain cell loss in PD may be preceded by their synaptic dysfunction and loss, and may subsequently be accompanied by tau phosphorylation and increases in insoluble α -synuclein.^44,47^ Therefore, in VKI mice treated with LPS, we examined microglial activation, phagocytic capacity and synaptic pruning in dorsolateral striatum where DA neuronal projections reside. In VKI mice we have previously described deficits in striatal dopaminergic neurotransmission with amphetamine-induced hyperlocomotion, which are rescued by Lrrk2 kinase inhibition, and prior to any movement disorder resulting from neurodegeneration.^30^ Microglial activation in response to LPS is evident as their somas are enlarged (Fig. 6a, c) and irregular (Fig. 6d), and this phenomenon is further potentiated in VKI mice as illustrated by hypertrophic filament length (Fig. 6e) and Sholl intersections (Fig. 6f). IHC revealed LPS elevated microglial expression of Cd68, with no discrete genotypic differences (Fig. 6g). Moreover, LPS significantly elevated LPL (Fig. 6b, h). In contrast, VKI mice had a noticeable increase in baseline microglial LPL relative to WT, and upregulate LPL levels in response to LPS, although the amplitude of expression is markedly diminished compared to LPS-treated WT mice (Fig. 6h). These findings together with the scRNAseq dataset suggest altered lipid metabolism in naive VKI mice, that through adaptation has established a suppressive phenotype to mitigate this response. Microglia constantly survey the brain parenchyma, even pruning synapses under homeostatic conditions. LPS induced inflammation elevated microglial pruning of synapses as shown by the detection of internalized bassoon and homer1, pan-markers of pre-and post-synapses, respectively. In LPS treated mice, Vps35 p.D620N appears to exacerbate this process, as evident by increased bassoon immunoreactivity within Iba1+ microglia (Fig. 7a–c), despite further phagocytotic activity with LPS being blunted.

**Fig. 6.**
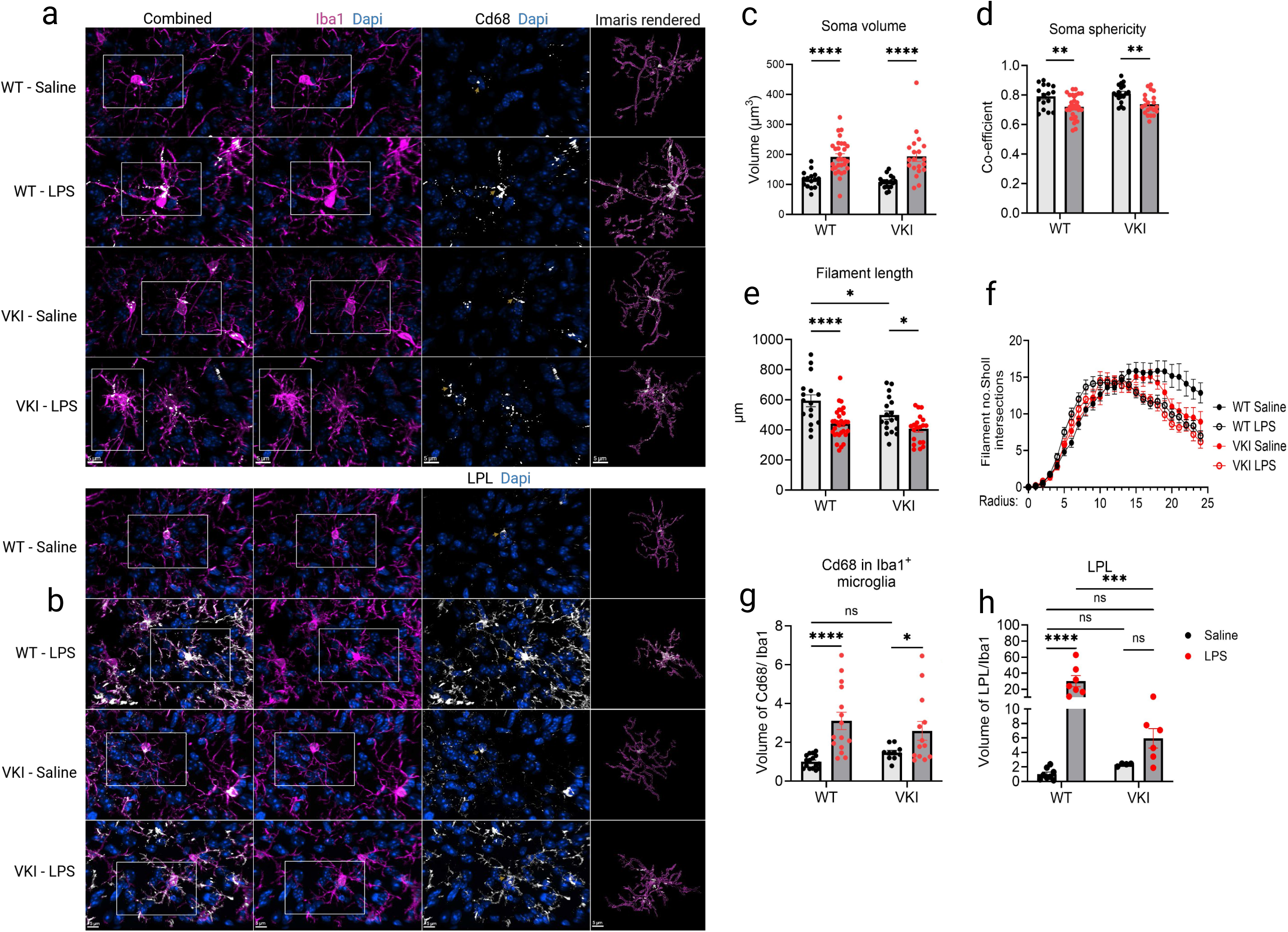
Vps35 p.D620N exacerbates microglia activation and phagocytosis under inflammatory conditions in the striatum. a, **b)** IF-CM representative images for Iba1, Cd68 and LPL co-stained with Dapi, and their Imaris^TM^ rendering in striatum of WT and VKI^Hom^ mice, with/without LPS. Two-way ANOVA with Tukey’s multiple comparisons test of: microglial soma volume (**c**, WT vs WT LPS, ****p<0.0001; VKI vs VKI LPS,****p<0.0001), microglial soma sphericity (**d**, WT vs WT LPS, **p = 0.0029; VKI vs VKI LPS, **p = 0.0028) and microglial filament length (**e** WT vs WT LPS, ****p<0.0001; WT vs VKI, *p = 0.0195; VKI vs VKI LPS, *p = 0.0184). **f**) Number of filaments Sholl intersections: WT n = 2 (16); WT LPS n = 3 (30); VKI^Hom^ n = 2 (18) VKI^Hom^ LPS = 2 (20). **g)** Two-way ANOVA with Tukey’s multiple comparisons test of the volume of Cd68+ puncta in Iba1^+^ microglia (WT vs WT LPS, ****p<0.0001; VKI vs VKI LPS, *p = 0.0417). [WT n = 4 (8); WT LPS n = 6 (11); VKI^Hom^ n = 3 (6); VKI^Hom^ LPS = 4 (8)]. **h)** Two-way ANOVA with Tukey’s multiple comparisons test of the volume of LPL+ puncta in Iba1^+^ microglia (WT vs WT LPS, ****p<0.0001; WT LPS vs VKI LPS, ***p = 0.0002). [WT n = 4 (8); WT LPS n = 4 (7); VKI^Hom^ n = 3(5); VKI^Hom^ LPS = 3(6)]. *p <0.05, **p<0.01, *\*\*\*p*<0.001, \**\*\*\*p*<0.0001. Error bars are ± SEM. n refers to the number of mice and in parentheses the number of tissue sections or number of microglia analyzed for Fig. 6c - f. The yellow arrow denotes the punctas.

**Fig. 7.**
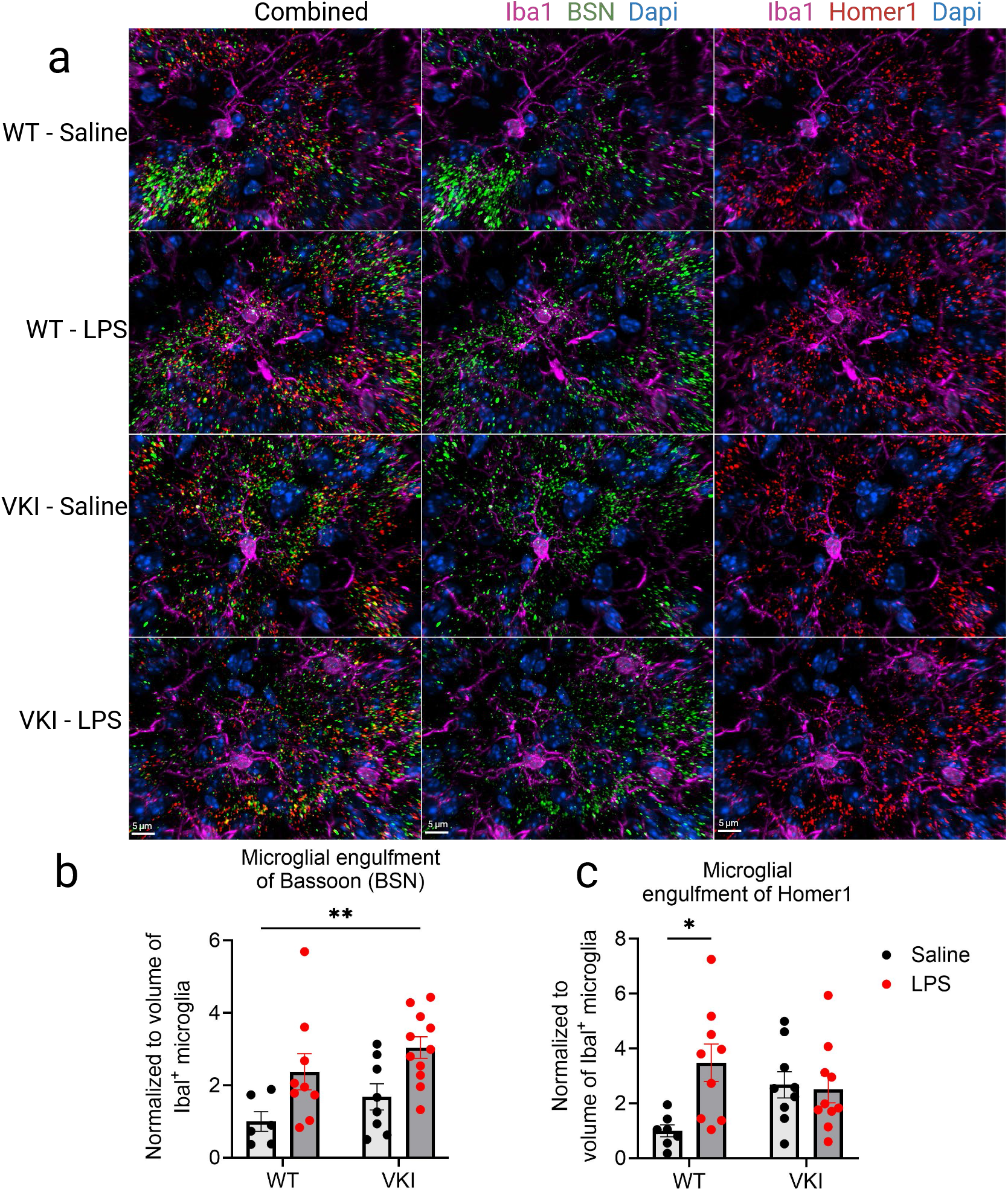
Vps35 p.D620N may increase vulnerability to microglia mediated-synaptic pruning. **a)** IF-CM images for Iba1, Bassoon (BSN) and Homer1 co-stained with Dapi in the striatum of WT and VKI^Hom^ mice, with/without LPS. Two-way ANOVA with Tukey’s multiple comparisons test of: BSN^+^ puncta in Iba1^+^ microglia (**b**, WT vs WT LPS, p = 0.1446; VKI vs VKI LPS, p = 0.0759; WT vs VKI LPS, **p = 0.0064) and Homer1^+^ puncta in Iba1^+^ microglia (WT vs WT LPS, *p = 0.0167). [WT n = 3 (6); WT LPS n = 5 (9); VKI^Hom^ n = 3(9); VKI^Hom^ LPS = 4(11)]. *p <0.05, **p<0.01. Error bars are ±SEM. n refers to the number of mice and in parentheses the number of tissue sections.

## DISCUSSION

*LRRK2* gene and promoter polymorphisms are modestly associated with PD and several chronic immune and inflammatory disorders^7,48^, and *LRRK2* expression is immunoresponsive in immune cells. Nevertheless, *LRRK2* mutations genetically linked to PD, that are causal and not merely associated, all increase endogenous LRRK2 kinase activity. Among known variants in *LRRK2, RAB 32 and VPS 35 , in vitro/ex vivo* studies demonstrate VPS35 p.D620N increases LRRK2 kinase activity the most.^23,49^ Hence, this study aimed to explore transcriptomic and functional consequences of Vps35 p.D620N on microglia in VKI mice, together with the effects of peripheral immune stimulation. We reveal that Vps35 p.D620N drives microglia into a chronic, partially reprogrammed state characterized by coordinated upregulation of humoral antimicrobial/alarmin genes (e.g. *S100a8*, *S100a9*, *Lcn2*, *Camp*, *Ngp*, *Ltf* and *Retnlg*) with concomitant down regulation of homeostatic environment sensing genes (*Adgrg5*, *Ccr2*, *Gpr68*). At baseline, VKI microglia are morphologically modestly hypertrophic, a phenomenon exacerbated with LPS induced inflammation, which concomitantly suppressed microglial Trem2. These findings were verified in cVKI mice that express 50% of Vps35 protein. Peripheral inflammation with LPS in both WT and VKI activates brain microglia to increase striatal synaptic engulfment, and Vps35 p.D620N synergizes that effect. LPS also increases microglial LPL in WT, but this response is blunted in VKI striatum, which reveals a deficit in their inducible lipid-handling machinery despite preserved activation capacity.

WT and VKI mice were singly-sexed, group housed in sterilized microisolator caging, using sterilized bedding, chow, and HEPA-filtered air, and consequently had minimal and comparable exogenous pathogen exposure. However, scRNAseq on enriched microglia extracted from the brain of VKI mice, revealed that Vps35 p.D620N chronically upregulates genes involved in pathogen defense, innate immune responses and phagocytosis, and those results are directly attributable to animal genotype. Alarmin genes are classical markers of neutrophils which may evoke skepticism for their microglial origin. However, we identified a distinct, independent, neutrophil cluster in our dataset, and we have validated alarmin protein expression in microglia, including S100a9 and Lcn2 by IHC. These complimentary findings provide assurance that microglial scRNAseq expression of alarmin genes is not a consequence of neutrophil contamination. The VPS35 p.D620 residue is a mutational hotspot — a structural or chemically vulnerable site at which pathogenic mutations recur, albeit on different haplotypes. Although this mechanism contrasts to the most frequent, pathogenic mutations in *LRRK2* and *RAB32*, that are inherited ‘identical-by-descent’ due to positive selection, it is also indicative of an evolutionary advantage to resist pathogen-mediated retromer subversion.^4^ Notably, *Legionella pneumophila*, *chlamydia trachomatis*, human immunodeficiency virus (HIV), and several other intracellular pathogens encode effectors that directly target retromer components to redirect endosomal trafficking and establish replicative niches.^50^ VPS35 p.D620N may reflect evolutionary pressure at the precise residue where host-pathogen interaction shapes retromer function, consistent with antagonistic pleiotropy.^4,51,52^ The age-associated consequences of chronic microglial activation may contribute to the insidious demise of DA neurons in *VPS35* Parkinson’s disease.^53^ Nevertheless, careful assessment of microglia subtypes, and the transcriptomic repertoires in our dataset reveal no change in the proportion of interferon-responsive microglia, translationally primed/activated microglia, and DAMs. Rather, we identified a unique microglial subpopulation similar to DAMs but enriched with S100 calcium binding protein family genes (*S100a8* and *S100a9*), concomitant with high expression of *H2-K1*, *Apoe*, *Cst7* and *Lpl*. Our findings are complimentary to Gruel and colleagues who identified a *S100a8* and *S100a9* microglia subpopulation in 9-month-old senescence accelerated mouse-prone 8 (SAMP8), and *MAPT* p.P301L (tau) transgenic mice (K18-seeded P301L).^54,55^ The S100a8-S100a9 complex appears to contribute to amyloid pathology and cognitive deficits in mouse models of AD, and complementary human studies report increased microglial S100a8 in AD brain.^54^ Collectively, these studies verify the S100a8-S100a9 enriched microglia population in our data is legitimate and disease-relevant. Interestingly, unlike SAMP8 and tau transgenic mice, VKI at 6M exhibit no overt protein aggregates or aging phenotype, yet this microglial population emerges as a function of a single mutation. VKI mice are a physiologically faithful model of gene dysfunction in PD, that exhibit disease-relevant neuro-modulatory, behavioral and inflammatory phenotypes, without induction of hyperphysiologic conditions.

Importantly, the S100 family genes together with the other alarmin genes identified in our dataset constitute the senescence-associated secretory phenotype (SASP). These secretory humoral immune antimicrobial peptides can modify their immediate surroundings, propagating/enhancing the immune response.^52,56,57^ Not only is this response reminiscent of WT mice treated with peripheral LPS, but microglial activation is further potentiated by LPS treatment on a Vps35 p.D620N background, highlighting the genetic predisposition to inflammation. SASP plays a crucial role in senescence transmission through paracrine signaling to neighboring cells, creating a chronic, low- grade inflammatory environment, known as inflammaging. SASP not only recruits immune cells to eliminate senescent cells, but it also contributes to inflammation and tissue remodeling. The latter is exemplified physiologically as VKI have modestly increased vulnerability to microglial synaptic pruning. LPS induces phagocytosis and the formation of lipid droplets that increased levels of LPL are juxtaposed to breakdown.^58^ Hence, LPS robustly upregulates the expression of LPL in WT compared to their saline counterparts but, in contrast, LPL levels and phagocytotic responses in VKI striatum are blunted under inflammatory conditions. The striatal LPL phenotype is reminiscent of lipid-droplet associated microglia (LDAM), a novel microglia state identified in aged mice by Marschallinger and colleagues with a unique transcriptomic signature, phagocytic deficits and pro-inflammatory phenotype.^58^ These authors identified Vps35 as a negative regulator of lipid droplet formation in BV2 cells, a murine model of microglia. Their findings are congruent with LPS-induced inflammation in VKI mice and the blunted LPL levels we observe *in vivo*. In turn, increased lipid droplets enhance cytokine and chemokine levels, amplifying the SASP phenotype. These findings in VKI striatum are contrary to midbrain observations made in parallel, mood disorders are an early and prodromal feature of PD and more severe striatal involvement, disrupting frontostriatal reward and decision-making logic, may contribute to the high suicide rate in some VPS35 p.D620N families.^53^ Interestingly, Trem2 is downregulated by pro-inflammatory cytokines, the constituents of SASP as reported in microglial cultures^59,60^, although the mechanism remains elusive. This receptor has a pivotal anti-inflammatory role^33^, and appears to modulate the transitions of DAMs.^15^ Considering that naïve VKI mice have a SASP phenotype at baseline, it is congruent they have greater suppression of microglial Trem2, along with exacerbated inflammation compared to WT with LPS. Therefore, modulating the levels of Trem2 may be beneficial to control inflammation, and its deleterious effects.

The *SNpc* is regionally enriched in microglia; however, despite the availability of this knock-in model, it has yet to be tested whether their SASP develops into a detectable microglia senescence. The cellular and temporal changes that concur with the insidious, but selective, demise of DA neurons can now be deduced. Further investigations in VKI mice should decipher whether DAMs-like microglia in our current dataset, with age, transition into bono fide DAMs to catalyze the severity of the disease, and might be extended to quantify nigral gliosis more broadly. Furthermore, the discrete role of LRRK2 kinase hyperactivity on microglial transcriptomic profile and function should be assessed, using specific inhibitors, to distinguish the former from retromer-specific effects on cargo trafficking. These experiments may also assess the long-term efficacy of Lrrk2 kinase inhibitors on neurodegeneration.

VPS35, LRRK2, RAB32 and RAB29 physically interact, and are intimately involved in endolysosomal cargo sorting and recycling versus lysosomal degradation. However, several major questions remain – how does VPS35 p.D620N induce a disproportionally high level of LRRK2 kinase activation? Lrrk2 kinase activity has been reported in iPSC-derived macrophages and microglia to suppress their degradative capacity^61^; thus, an answer to this question may shed light on some of the microglial specific deficits in VKI mice. We hypothesize Mendelian gene mutations merely represent molecular extremes of the same continuum perturbed in idiopathic Parkinson’s disease; minimally VPS35, RAB32 and LRRK2 converge on the same mechanism.^2,11^ Hence, pathophysiologic studies using these models, that encompass non-autonomous and dopaminergic cell-autonomous responses, may explain the selective vulnerability of the *SNpc*, its gliosis and the subsequent development of neuronal inclusion pathologies. Human *post-mortem* studies are complementary but are limited to end-stage. Physiological knock-in/knock-out models based on Mendelian genetic etiology not only provide mechanistic targets, but they also provide an orthogonal means to test this biology, as it develops, and relevant interventions to break this cycle, to enable neuroprotective and disease-modifying therapeutics.

## METHODS

### Mice engineering and husbandry

Vps35 constitutive knock-in (VKI) mice were engineered as previously described.^29,30^ Briefly, a cre-conditional Vps35 p.D620N knock-in (VKI; B6.Cg-*Vps35^tm1.1Mjff^*/J; Jax strain#023409) mouse model was developed via homologous recombination. We introduced a cDNA cassette of murine ‘exons 15-17, heterologous stop and polyA sequence’, immediately flanked by a pair of lox P sites. The 5’ vector arm is complementary to intron 14. The 3’ vector arm included intron 14 - exon 15, and the exon 15-point mutation encoding Vps35 p.D620N. The mini-gene insertion inadvertently silenced the expression of the recombinant allele to create haploinsufficient mice (cVKI) (50% reduction in Vps35 protein levels). The entire cDNA cassette was subsequently removed by breeding to germline Cre transgenic animals to just leave the constitutive knock in, VKI ((Vps35 g.85,263,520G>A (p.D620N) mutation (GRCm38/mm10; NM_022997.5)). Both strains are breeding in our vivarium at the University of Florida (UF), and are available through Jackson Laboratories. They have been maintained on the same C57BL/6J background [Jax strain #000664] for >10 generations. Animal studies were approved by the Institutional Animal Care and Use Committee at UF. All mice were kept on a reverse cycle (light on from 8:30 pm to 8:30 am) and single-sex group-housed in enrichment cages after weaning at post-natal day 21, in accordance with NIH guidelines for care and use of animals. For all experiments, 5–6M old male animals were used. Ear notches were taken for DNA extraction and genotype validation.

### Single cell RNA sequencing (scRNAseq)

5–6M old male VKI mice and their controls (WT n = 2, VKI^Het^ n = 3 and VKI^Hom^ n = 3) were anesthetized via inhalation of isoflurane, and transcardially perfused with PBS before brain collection. The brains were placed on a petri dish and cut into eight sagittal pieces using a scalpel then transferred into gentleMACS C-Tube containing digestion enzymes. Subsequently, we used the Miltenyi™ gentle mechanical dissociation method followed by CD11b^+^ antibody-based selection (mouse CD11b^+^ selection kit II, STEMCELL Technologies) to isolate microglia using published methods.^62^ Isolated microglia were counted, and their viability assessed before being subjected to 10x Genomics scRNAseq. Sequencing libraries were prepared following a published Smart-seq2 protocol and sequenced on an Illumina NovaSeq 6000.^63^ The number of total reads and genes detected were evaluated to filter out cells with low sequencing quality.^36^ We used the Seurat v4 package to perform unsupervised clustering analysis on scRNAseq data.^36,63^ Sequence reads from the Illumina NovaSeq 6000 platform were processed using the Cell Ranger (v9.0.0) with the mouse genome (mm10) as the reference genome. After the initial QC (barcoding filtering, UMI counting), the filtered matrices were imported to Seurat package (v5.4.0) in R (v4.4) for analysis. In Seurat, the PCA and UMAP were used for dimensionality reduction, and the clustering was performed using FindNeighbors and FindClusters functions. The Clustree R package (v0.5.1) was used for analyzing clustering results. Cell type annotation was performed using the Cellmarkeraccordion R package (v1.0.0) based on the database of CellMarker2.0. Differential expression analysis was conducted with the FindMarkers function in Seurat and the results are presented in a volcano plot generated with the EnhancedVolcano R package (v1.22.0). Customized gene markers in clusters were visualized in dimensional reduction plots generated with the Feature Plot function of Seurat. Differentially expressed genes, defined as those with an adjusted p-value < 0.05 and fold change > 1.4, were separated into upregulated and downregulated gene sets for each comparison and analyzed independently using STRING for protein-protein interaction network generation and functional enrichment analysis. Gene symbols were entered together with their corresponding fold-change values, which were used to rank proteins and map the magnitude and direction of expression changes across networks. STRING analysis was performed using the full network, evidence-based edges, all available interaction sources, and a minimum interaction score of 0.400 (medium confidence), with disconnected nodes excluded from network display.^64^ Complimentary to the STRING analysis, we carried out pathway enrichment analysis (PEA) in g:Profiler g:GOSt (version: e113_eg59_p19_6be52918) via the g:SCS threshold methods at a p-value threshold of 0.05.^65^

### Immunohistochemistry (IHC)

For imaging, mice were anesthetized via inhalation of isoflurane, intracardially perfused with PBS and their brains were rapidly extracted then post-fixed overnight at 4°C in 4% paraformaldehyde (PFA). The brains were then cryoprotected in 30% sucrose in PBS before sagittal slices (30 µm) were obtained with a cryostat. Sections were blocked in 10% normal goat serum (NGS) or normal donkey serum (NDS) in 0.3% triton-X 100 in PBS at 37°C for 1 hour (h). However, for successful detection, certain antibodies required antigen retrieval which was carried out at 37^0^C for 1h in 1x tris-EDTA antigen retrieval buffer (Proteintech, Cat no: PR30002), then blocking with 0.3% glycine, 1% bovine serum albumin (Lot: 64758436) and 10% NGS. The sections were then incubated with antibodies of interest overnight at 4°C (**Supplemental Table 1**), which was followed by 4 washes in PBS, each 15 min, and secondary antibody incubation with goat or donkey species-specific Alexafluor IgG secondary antibodies (1h at RT, Invitrogen; 1:500). Tissues were rewashed in PBS (4x 15 min), then co-stained with DAPI (ThermoFisher Scientific,1:5000) and mounted using Fluoromount-G^TM^ (00-4958-02, Invitrogen©). Images were acquired at 40x or 60x field of view on a Nikon® Spinning Disc Confocal Microscope (AX/AX R with NSPARC) using NIS-Elements AR Imaging software from Nikon® (90 z- stack steps at 0.30 μm spacing), with its intrinsic deconvolution function. Immunoreactivity in the midbrain and striatum was quantified with Imaris^TM^ software (Oxford Instruments). Briefly, Iba1^+^ microglia (smoothing; 0.10 μm) were rendered as surfaces, then identified using the AI component of the Imaris^TM^, with only whole cells in the field of view included in the analysis as identified with DAPI. The microglia projections that broke off from their somas were stitched before the total volume was quantified accordingly. Similarly, all the other proteins of interest were rendered as surfaces with smoothing factors of 0.10 μm at 10 voxels but rendered by their absolute intensity which varied according to the stain. The shortest distance to surface filter component in Imaris^TM^ was used to target microglial specific expression of the protein of interest. All the given volumes were quantified accordingly and normalized to total volume of Iba1. For Sholl analysis, microglia were rendered as filaments.^41,66^

### Single strand complementary DNA (cDNA) synthesis and real quantitative PCR (RT-qPCR)

RNA was extracted from the midbrain of WT and VKI mice then converted into cDNA using High-Capacity cDNA Reverse Transcription Kit (Applied Biosystems™). Briefly, in a 384-well plate, a mixture containing 10 μL of total cDNA master mix (10x RT Buffer, 10x RT Random Primers, 25x dNTP Mix [100 mM], MultiScribe™ Reverse Transcriptase [50 U/μL]) and 10 μL of 400 ng of RNA (diluted with RNase-free water) were dispensed across three wells per mouse (3 technical replicates). The 384-well plate was then sealed with optical tape and placed in a VeritiPro™ 384-well Thermal Cycler (Applied Biosystems™) to undergo reverse transcription as instructed in the High-Capacity cDNA Reverse Transcription Kit manual. After cDNA conversion, 4 μL from each well were pipetted into a new 384-well plate, where 10 μL of TaqMan™ Universal Master Mix II, with UNG (Applied Biosystems™), 5 μL of RNase/DNase-free water, and 1 μL of TaqMan™ Gene Expression Assay (Thermo Fisher Scientific©) was added in three wells per mouse, per gene examined. TaqMan™ Gene expression assay included: *glyceraldehyde-3-phosophate dehydrogenase (Gapdh)*: Mm99999915_g1; *beta-actin (Actb)*: Mm02619580_g1; *ionized calcium binding adaptor molecule 1* (*Aif1/Iba1*): Mm00479862_g1; Cd*68*: Mm03047343_m1; *Lcn2*: Mm01324470_m1 and *S100a9*: Mm00656925_m1. Once cDNA and TaqMan™ master mixes were pipetted, the 384-well plates were placed into a QuantStudio™ 5 Real-Time PCR System (Applied Biosystems™), where QuantStudio™ Design & Analysis Software (Applied Biosystems™) was used to conduct qPCR and find comparative Ct values (ΔΔCt) from each well using the standard protocols supplied in the software. For each mouse examined on the qPCR plates, the combined mean of Ct values from the housekeeping genes *Gapdh* and *Actb* were calculated and ΔCt was found for each targeted gene from each sample (ΔCt = Ct value of targeted gene – Ct value of combined mean *Gapdh* and *Actb*). To compare groups, the ΔΔCt method was used, where ΔΔCt = ΔCt of targeted gene from experimental group – ΔCt of targeted gene from control group. Once ΔΔCt was found for each comparison, relative gene expression for the gene of interest in each sample was calculated using 2^(-ΔΔCt)^.

### Western blotting

VKI and cVKI mice were euthanized and perfused per our previous protocols.^66^ The midbrains were microdissected and lysed in a Dounce homogenizer with 1x Cell Signaling Lysis Buffer (#9803S). Denatured lysates were resolved by SDS-PAGE on 4-20% TGX Criterion TGX pre-cast gels (Bio-rad) and transferred to nitrocellulose membranes by semi-dry trans-blot turbo transfer system (Bio-rad). The membranes were blocked with either 50% Intercept tris buffered saline (TBS; 927-60001), 5% bovine serum albumin (BSA) in TBS-Tween (TBST), or 5% milk in TBST for 1h at room temperature (RT) followed by incubation overnight at 4°C with primary antibodies diluted in blocking buffers depending on the protein of interest (**Supplemental Table 1**). After incubation with primary antibodies, the membranes were washed in TBST (3 x 5 min), followed by incubation for 1h at RT with fluorescently conjugated goat anti-mouse or rabbit IR Dye 680 or 800 antibodies (Licor). The blots were washed in TBST (3x 5 min) and scanned on a ChemiDoc MP imaging platform (Bio-Rad).^30,66^

### Statistical Analysis

IHC images were acquired via NIS-Elements AR software and quantified with the Imaris^TM^ software (Oxford Instruments), and the densitometric analyses for the immunoblots were performed with Image J software. Normality testing and statistical analyses of all the data were performed using GraphPad Prism 10 (GraphPad Software, San Diego, CA). The data is presented as mean ± standard error of the mean (SEM).

## Author contribution

IBD: conceptualization and project design, data curation, formal analysis, writing original draft, review and editing; MB: conceptualization and project design, data curation, formal analysis, review and editing, funding; JF: data curation and formal analysis, review and editing; RS: data curation and formal analysis, review and editing; AM: formal analysis, review and editing; LX: project design, data curation and formal analysis; FY: data curation and formal analysis; GR: data curation; SW: data curation, editing and reviewing; MJF: resources and funding, conceptualization and project design, training and supervision, critical review, interpretation and editing.

## Data availability

The VKI and cVKI strains have been deposited in Jackson Labs ((www.jax.org; *Vps35* knock-in: B6(Cg)- Vps35tm1.1Mjff/J; Stock No: 023409; floxΔneo Vps35; B6.Cg-Vps35tm1.2Mjff/J; Stock No: 021807)) with open distribution supported by the Michael J. Fox Foundation. Note, Vsp35 floxΔneo cassette silences the integrated allele, so cVKI are unable to be bred to homozygosity. The datasets generated during the current study are available from the corresponding authors and will be shared upon request.

## Supplemental material

**Fig. S1.**
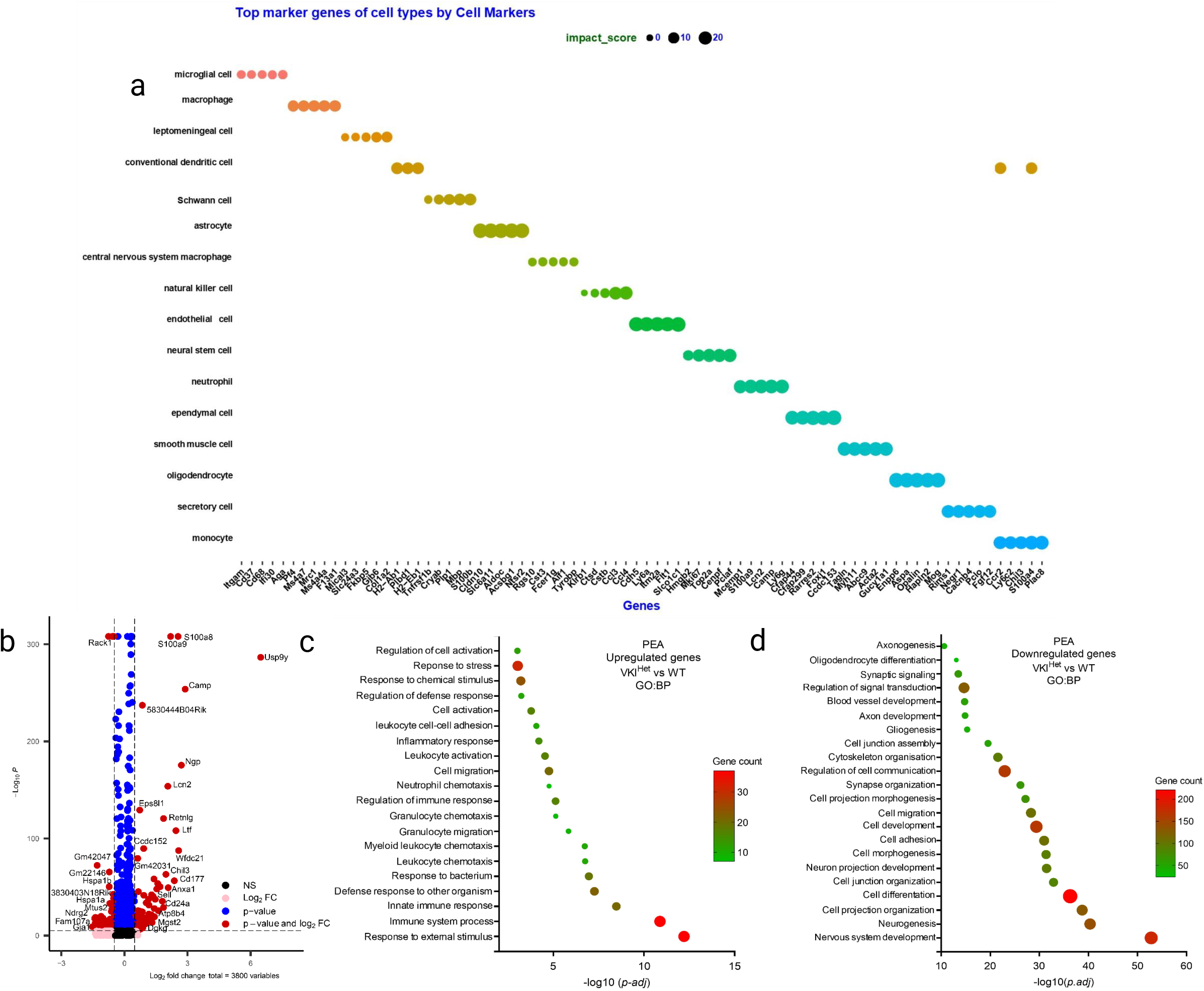
10x Genomics scRNAseq of CD11b^+^ antibody-based captured microglia from whole brain of VKI mice at ∼6M. **a**) Prediction of the different cell types based on their gene markers. **b**) Volcano plot generated with the EnhancedVolcano R package for upregulated and downregulated genes for all cell types in VKI^Het^. **c, d**) g:Profiler g:GOSt pathway enrichment analysis for upregulated and downregulated genes for all the cell types in VKI^Het^.

**Fig. S2.**
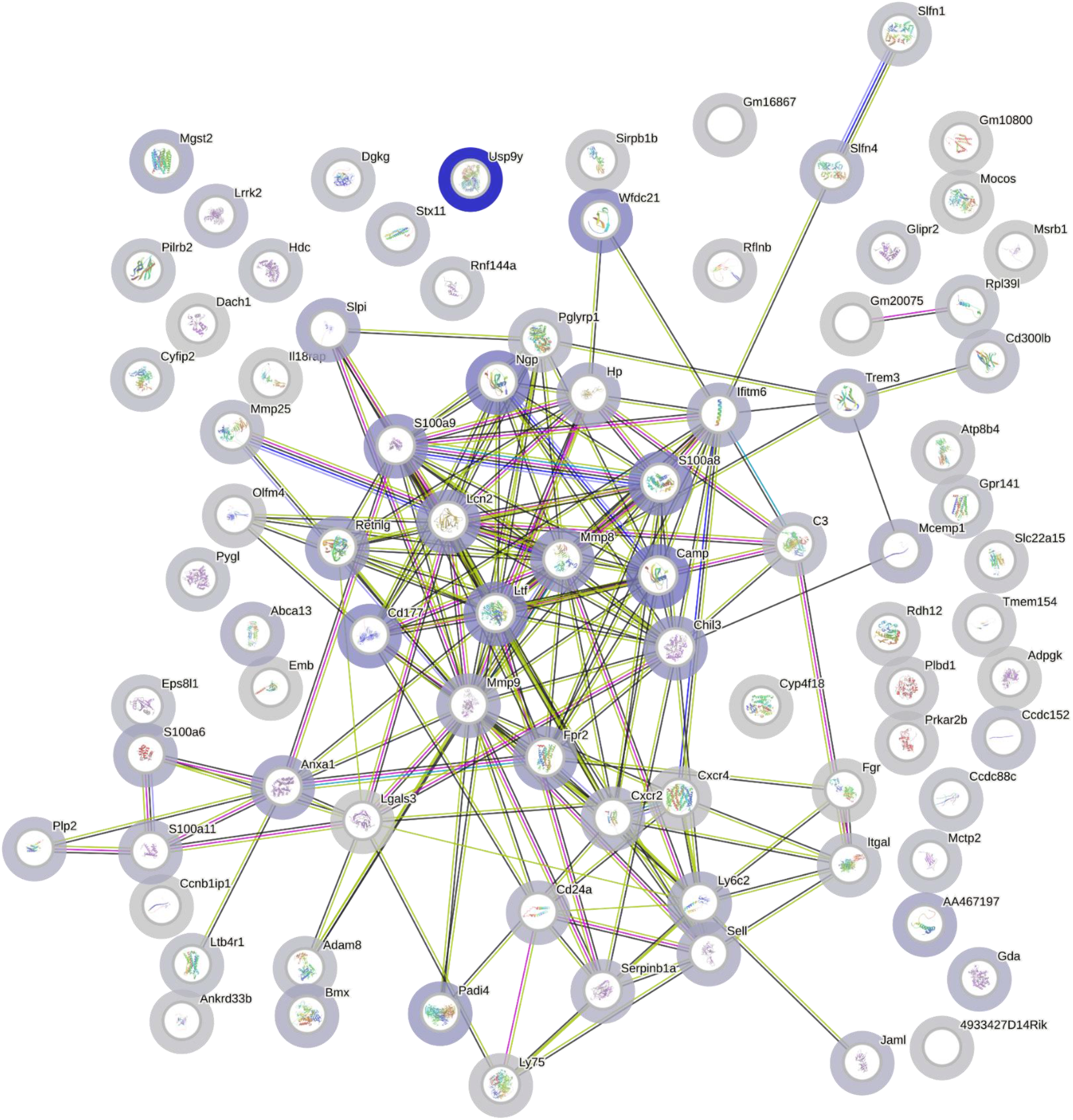
STRING generated protein-protein interaction network and functional enrichment analysis. VKI^Het^ Vs WT upregulated genes (from Fig. 1) **1. Core innate inflammatory module** •*S100a8*, *S100a9*, *Lcn2*, *Retnlg*, *Camp*, *Ltf* → Canonical myeloid inflammatory effector genes. •Suggests inflammatory activation as in the VKI^Hom^ **2. Complement / immune amplification** •*C3* → Central complement component. •Indicates early complement engagement even at the heterozygous stage. Not as broad as Hom, but clearly present. **3. Lysosomal / phagocytic remodeling** •*Lgals3*, *Anxa1*, *Mmp8*, *Mmp9*, *Chil3* → Phagolysosomal remodeling and matrix degradation genes. •Suggests emerging lysosomal stress and active tissue remodeling. This aligns with retromer (Vps35) dysfunction → lysosomal trafficking stress. **4. Chemokine / immune recruitment signaling** •*Cxcr2*, *Cxcr4*, *Ly6c2*, *Cd177*, *Sell* → Immune trafficking and recruitment-associated genes. •Suggests altered immune cell communication and possibly early immune cell recruitment. **5. Lipid handling / immune-metabolic shift** •*Hp*, *Abca13*, *Emb* → Acute-phase and lipid-associated genes. •Consistent with early metabolic reprogramming of activated microglia.

**Fig. S3.**
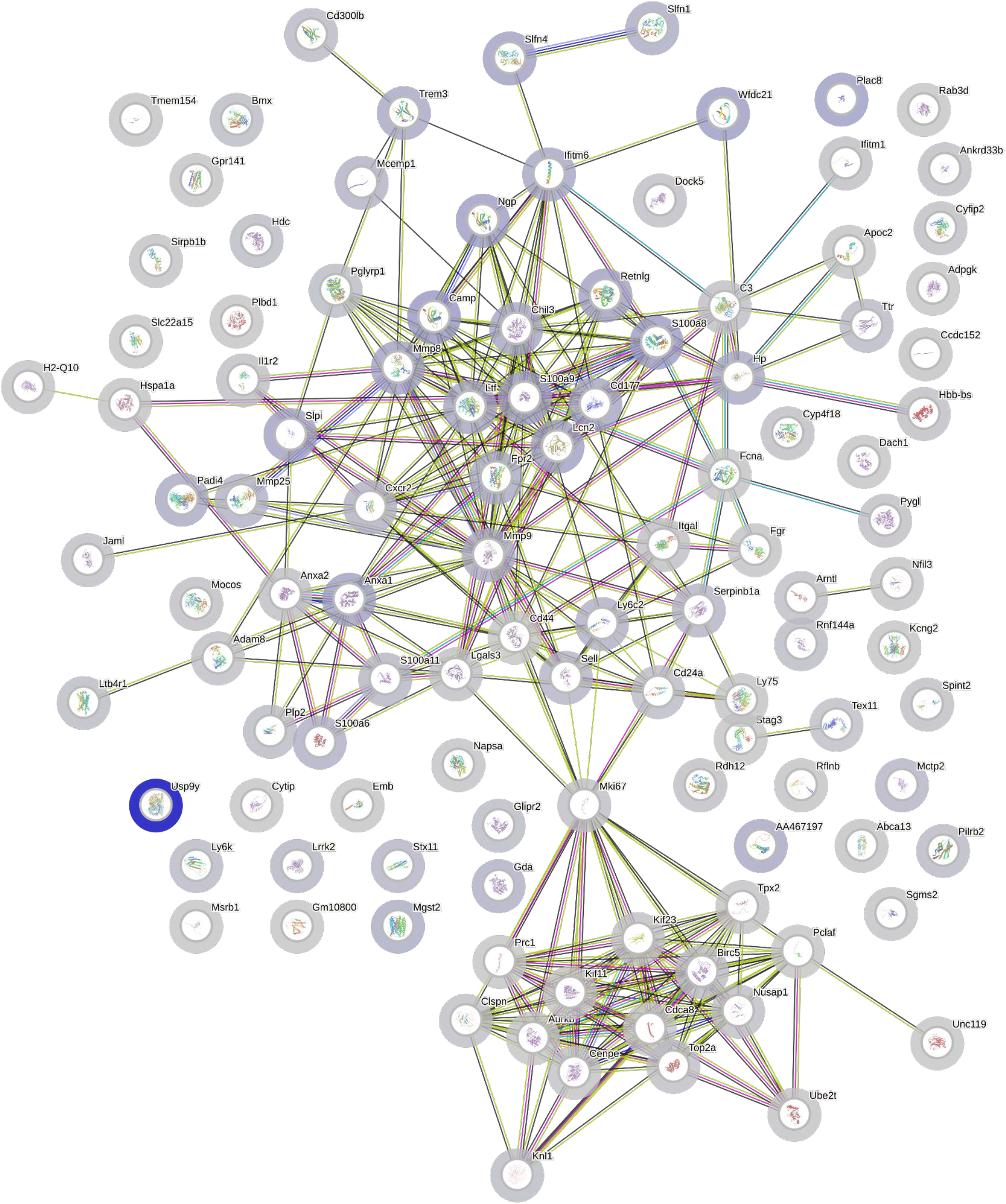
STRING generated protein-protein interaction network and functional enrichment analysis. VKI^Hom^ vs WT upregulated genes (From Fig. 1) 1. Acute inflammatory / NFκB – IL-1 axis cluster •*IL1b, Lcn2, S100a8, S100a9, Retnlg, Camp* and *Cxcl2* → Core innate inflammatory effector genes. •Suggests strong activation of the IL-1 / NF-κB pathway and a myeloid inflammatory program visible at the tissue level. **2. Complement / innate immune amplification** •*C3* and *Hp* → Complement activation and acute-phase response components. •Indicates propagation of inflammatory signaling beyond microglia, consistent with tissue-level immune remodeling. **3. Lysosomal stress / phagocytic activation** •*Lgals3, Anxa1, Anxa2, Trem3, Mmp9* and *Mmp8* → Phagolysosomal remodeling and extracellular matrix degradation. •Supports lysosomal stress and active phagocytic remodeling, coherent with VPS35-related endosomal trafficking defects. **4. Matrix remodeling / tissue remodeling** •*Mmp8, Mmp9, Itgal* and *Fgr* → ECM remodeling and immune-cell activation markers. Suggests structural reorganization accompanying inflammation. **5. Lipid handling / metabolic reprogramming** •*Apoe, Abca13* and *Serpinb1a* → Lipid transport and immune-metabolic shift genes. •Consistent with activated microglial metabolic reprogramming (DAM-like features). **6. Proliferation / cell-cycle super-cluster** •*Mki67, Aurkb, Top2a, Cenpe, Kif11, Kif23, Prc1, Ube2c* and *Birc5* → Canonical mitotic cell-cycle machinery. •Suggests active cellular proliferation — most likely proliferating activated microglia (reactive gliosis).

**Fig. S4.**
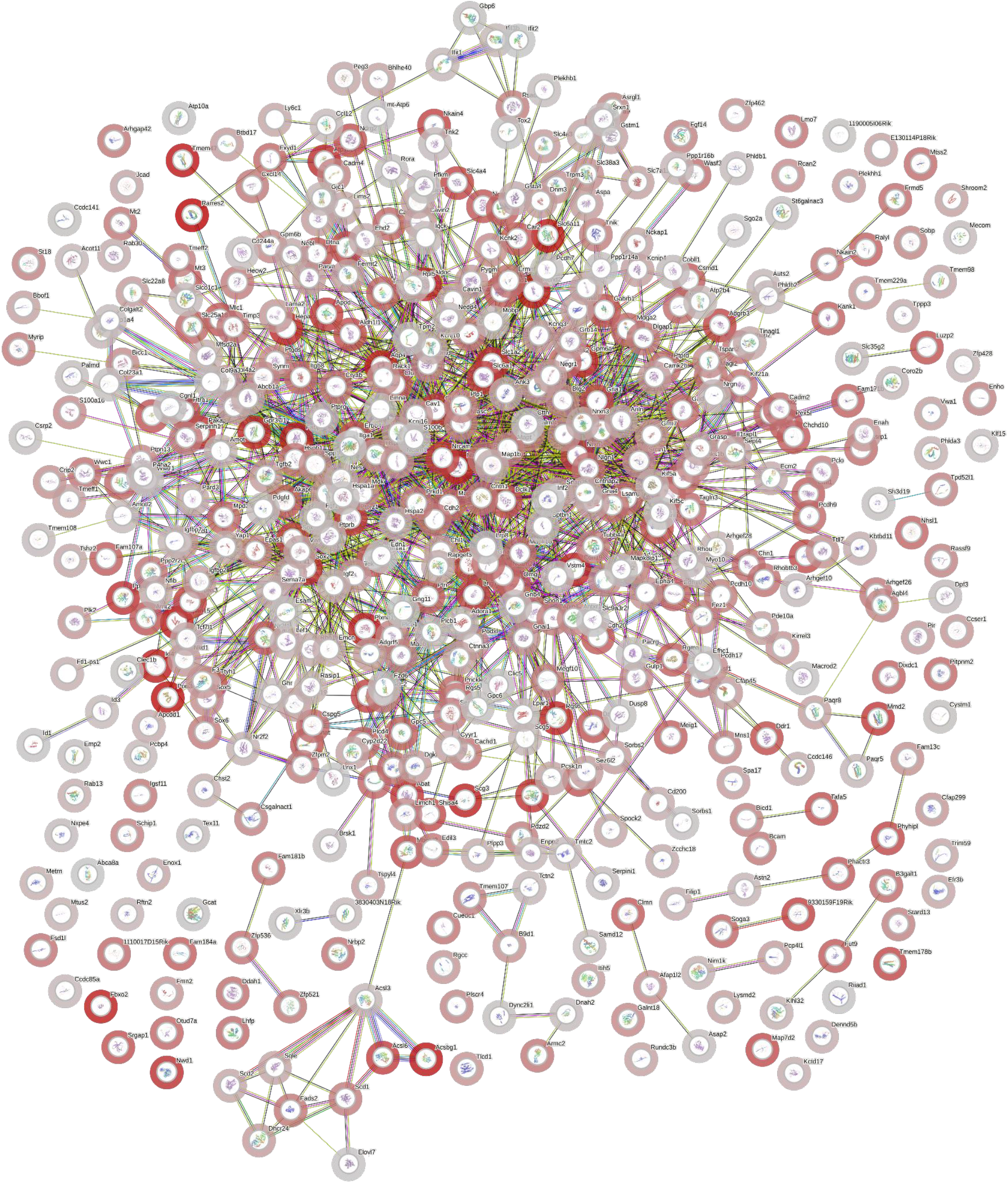
STRING generated protein-protein interaction network and functional enrichment analysis. VKI^Het^ Vs WT downregulated genes (from Fig. 1) 1. Synaptic transmission / neuronal signaling cluster •*Gria1*, *Gria2*, Grin family members, *Dlg4*, synaptic scaffolding genes → Glutamatergic synaptic machinery. •Suggests early suppression of excitatory synaptic programs. This mirrors VKI^Hom^ but appears less sharply clustered and less dominant. **2. Ion channel / neuronal excitability module** •Kcnj/Kcnq family members, Cacna-related genes → Voltage-gated channel components. •Indicates altered neuronal excitability gene expression, but likely subtler than in VKI^Hom^. **3. Axon guidance / neuronal adhesion** •*Nrcam, Ncam1*, related adhesion molecules → Neuronal structural integrity genes. •Suggests mild structural remodeling or altered neuron–glia interactions. **4. Synaptic vesicle / presynaptic machinery** •Presynaptic regulators and vesicle-associated genes (Snap/Syt-associated network nodes) → Neurotransmitter release machinery. •Suggests early synaptic dysfunction. **5. Metabolic / mitochondrial neuronal support genes** •Slc transporters and metabolic components scattered in network → Neuronal energy handling. •Suggests subtle metabolic stress in neurons.

**Fig. S5.**
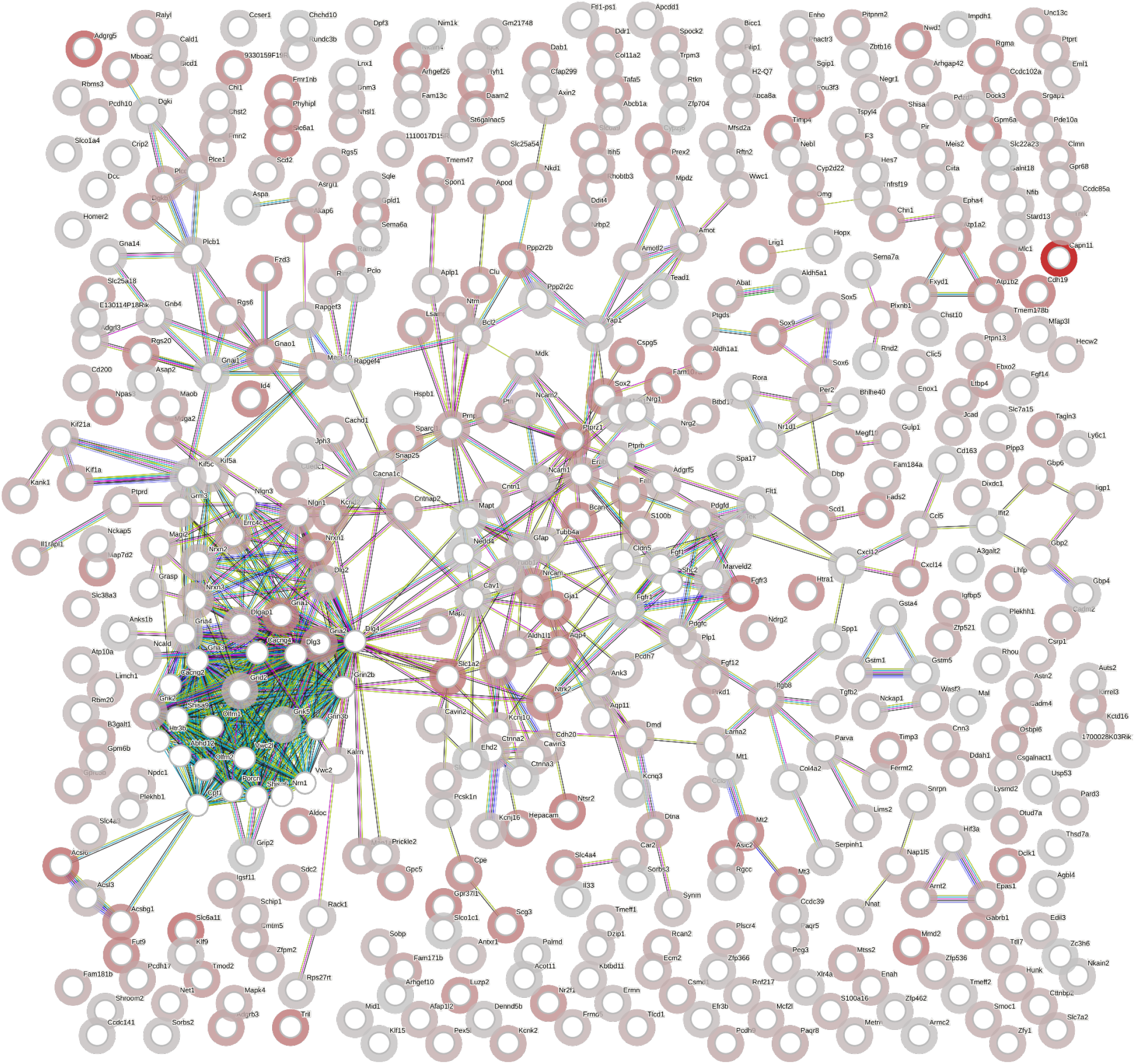
STRING generated protein-protein interaction network and functional enrichment analysis. VKI^Hom^ vs WT downregulated genes (from Fig. 1) 1. Synaptic transmission / neuronal signaling cluster •*Gria1, Gria2, Gria3, Grin1, Grin2b, Gabrb1, Gabra2, Dlg4* and *Shank*-associated genes → Glutamatergic and GABAergic synaptic components. •Suggests suppression of excitatory and inhibitory synaptic signaling at the tissue level. **2. Voltage-gated ion channel / excitability module** •*Kcnq3* and *Kcnj* family members, *Cacna1c*, Scn-related genes → Ion channel and membrane excitability components. •Indicates reduced neuronal electrical activity gene expression. **3. Axon guidance / structural neuronal maintenance** •*Nrcam, Ncam1 and L1cam*-associated components, cytoskeletal adaptors → Neuronal adhesion and axonal organization genes. •Suggests structural remodeling or neuronal stress. **4. Synaptic vesicle / neurotransmitter release machinery** •Synaptic vesicle trafficking genes (Snap/Syt/related regulators in network) → Presynaptic machinery. •Indicates impaired synaptic vesicle cycling or reduced synaptic density. **5. Neuronal metabolic support** •Slc family transporters (neuronal glucose/amino acid transporters), mitochondrial components → Neuronal metabolic support genes. •Suggests altered neuronal metabolic state.

**Fig. S6.**
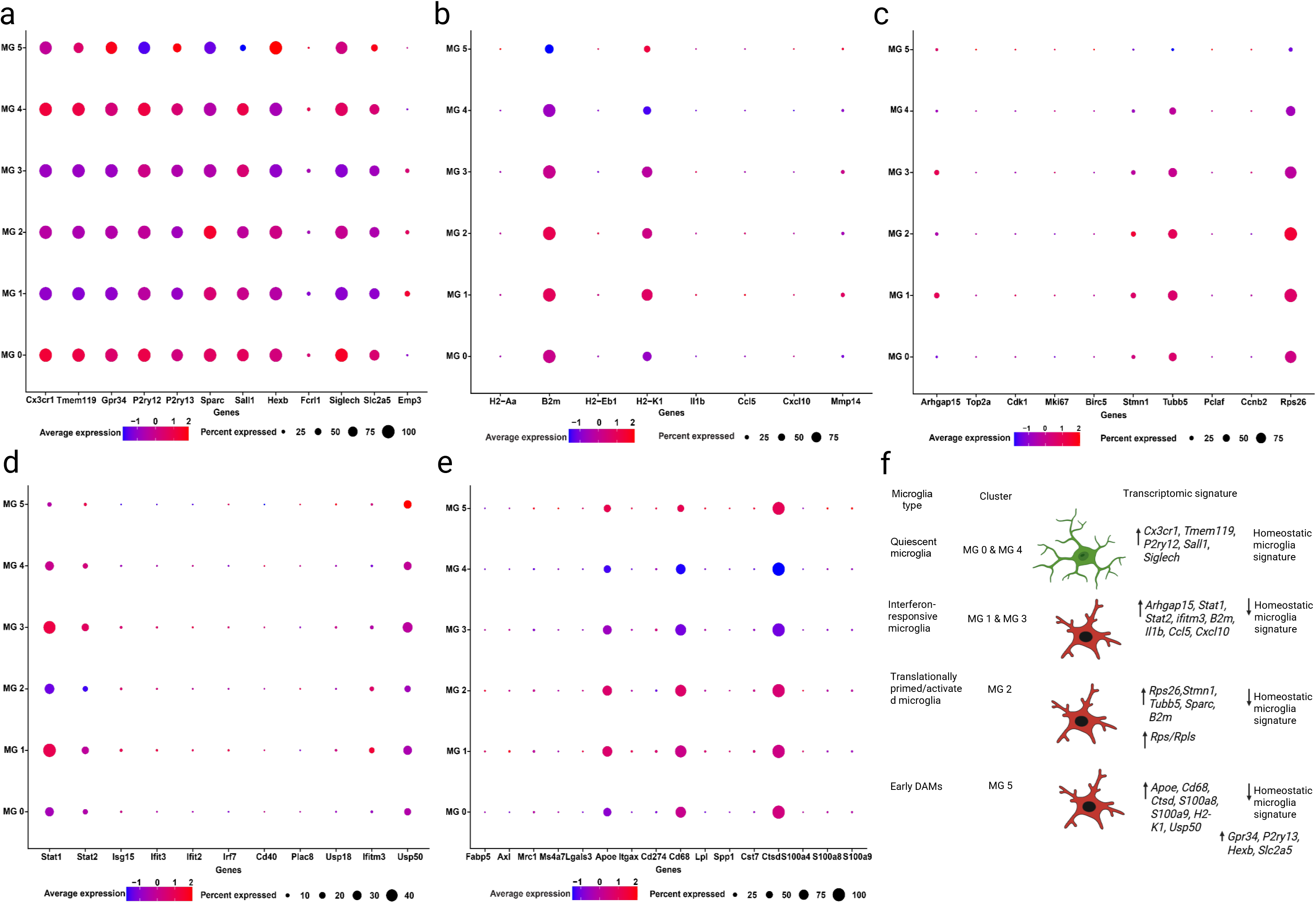
ScRNAseq revealed five microglia subtypes from CD11b^+^ antibody-based captured microglia from whole brain of VKI mice (both Het and Hom combined). For better visualization, the dot plot was segmented into five clusters as follows: **a)** enriched in homeostatic microglial genes, **b**) enriched in genes for antigen processing/presentation, and inflammatory/cytokine signaling, **c**) enriched in proliferation / cell-cycle genes, **d**) enriched in interferon-response genes, and **e**) enriched in DAMs genes. **f**) Classification of microglia-subtypes in VKI brains based on their transcriptomic signature.

**Fig. S7.**
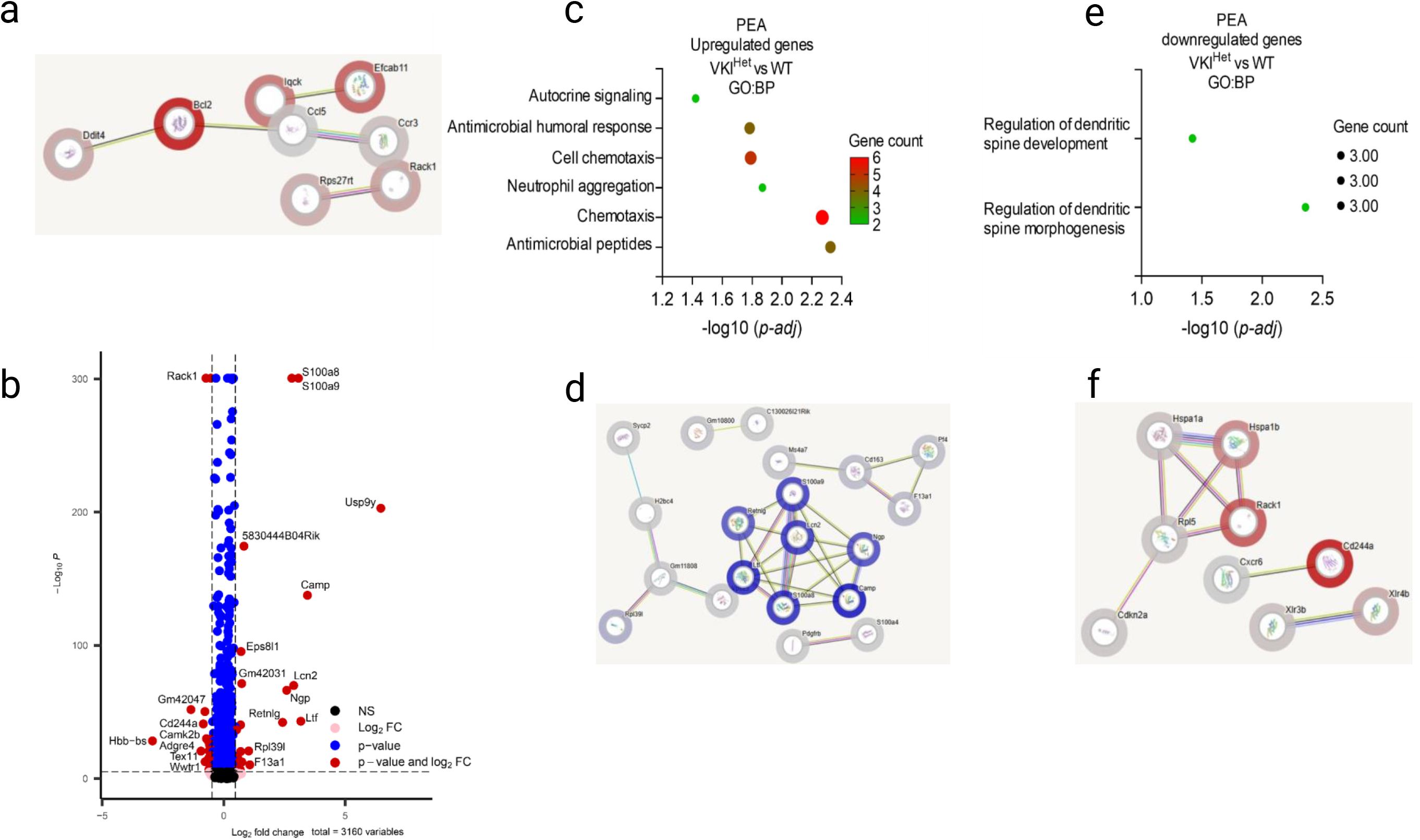
Vps35 p.D620N increases expression of microglial genes involved in innate immunity/acute inflammation, phagocytosis and lysosomal stress. **a**) STRING generated protein-protein interaction network and functional enrichment analysis for downregulated genes in VKI^Hom^ enriched microglia, with disconnected nodes excluded from network display. **b**) Volcano plot generated with the EnhancedVolcano R package for upregulated and downregulated genes in VKI^Het^ enriched microglia. **c**) g:Profiler g:GOSt pathway enrichment analysis for upregulated genes in VKI^Het^ microglia. **d**) STRING generated protein-protein interaction network and functional enrichment analysis for genes upregulated in VKI^Het^ microglia. **e**) g:Profiler g:GOSt pathway enrichment analysis for downregulated genes in VKI^Het^ microglia. **f**) STRING generated protein-protein interaction network and functional enrichment analysis for downregulated genes in VKI^Het^ enriched microglia.

**Fig. S8.**
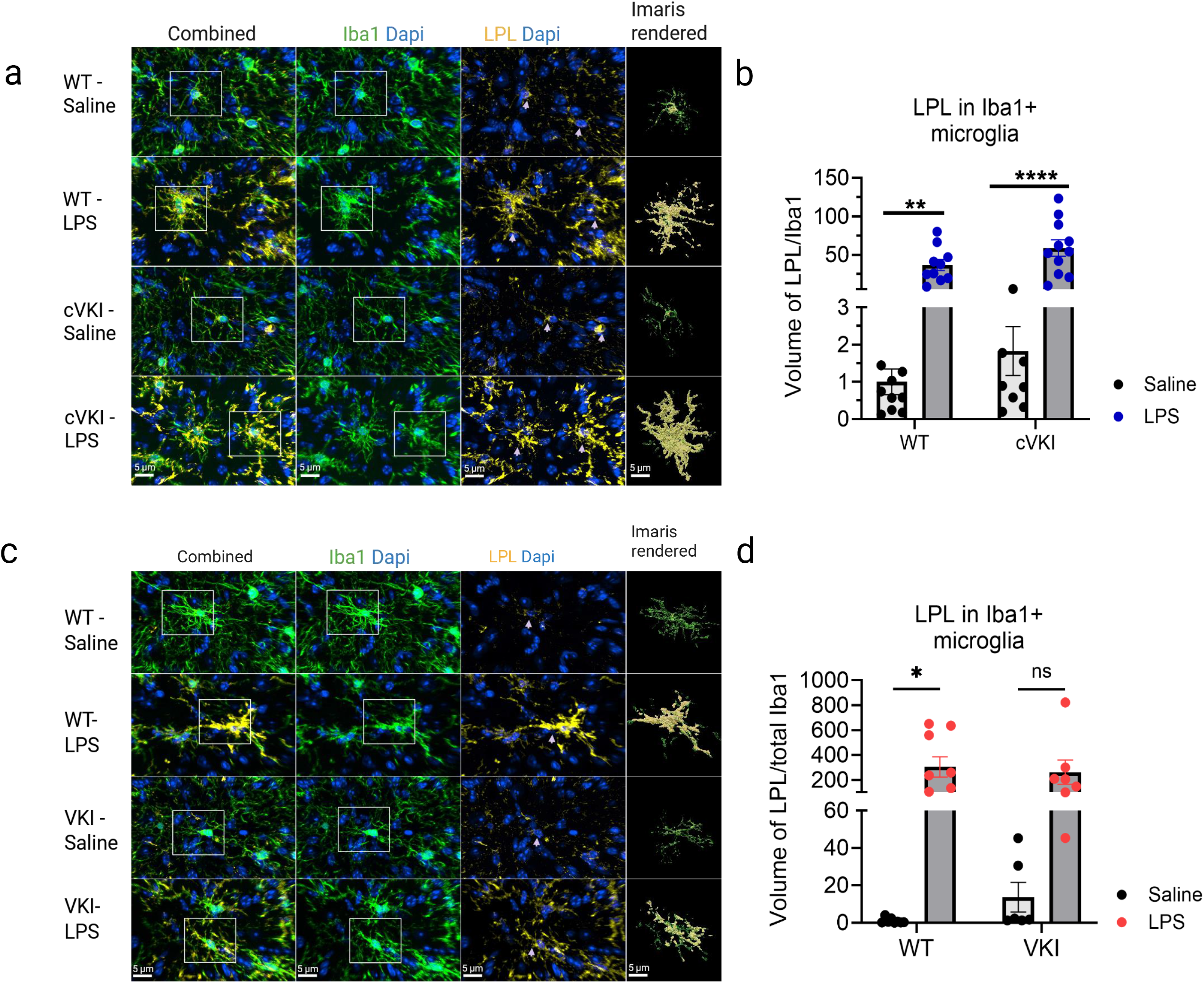
LPL expression is significantly elevated by systemic LPS, but no genotype dependent difference. **a)** Immunofluorescence confocal microscopy (IF-CM) representative images for Iba1 and LPL co-stained with Dapi, and their Imaris^TM^ rendering in the SN of WT and cVKI mice injected with LPS. **b**) Two-way ANOVA with Tukey’s multiple comparisons test of LPL immunoreactivity in Iba1+ microglia (WT vs WT LPS, **p = 0.0041; cVKI vs cVKI LPS, ****p<0.0001). [WT n = 5(10); WT LPS n = 5(10); cVKI n = 5(9); cVKI LPS = 6(11)]. **c)** IF-CM representative images for Iba1 and LPL co-stained with Dapi, and their Imaris^TM^ rendering in SN of WT and VKI^Hom^ mice injected with LPS. **d**) Two-way ANOVA with Tukey’s multiple comparisons test of LPL immunoreactivity in Iba1+ microglia (WT vs WT LPS, *p = 0.0109). [WT saline n = 4 (8); WT LPS n = 5 (9); VKI^Hom^ Saline n = 3(6); VKI^Hom^ LPS = 4(7)]. Error bars are ± SEM. n refers to the number of mice and in parentheses the number of sections.

**Fig. S9.**
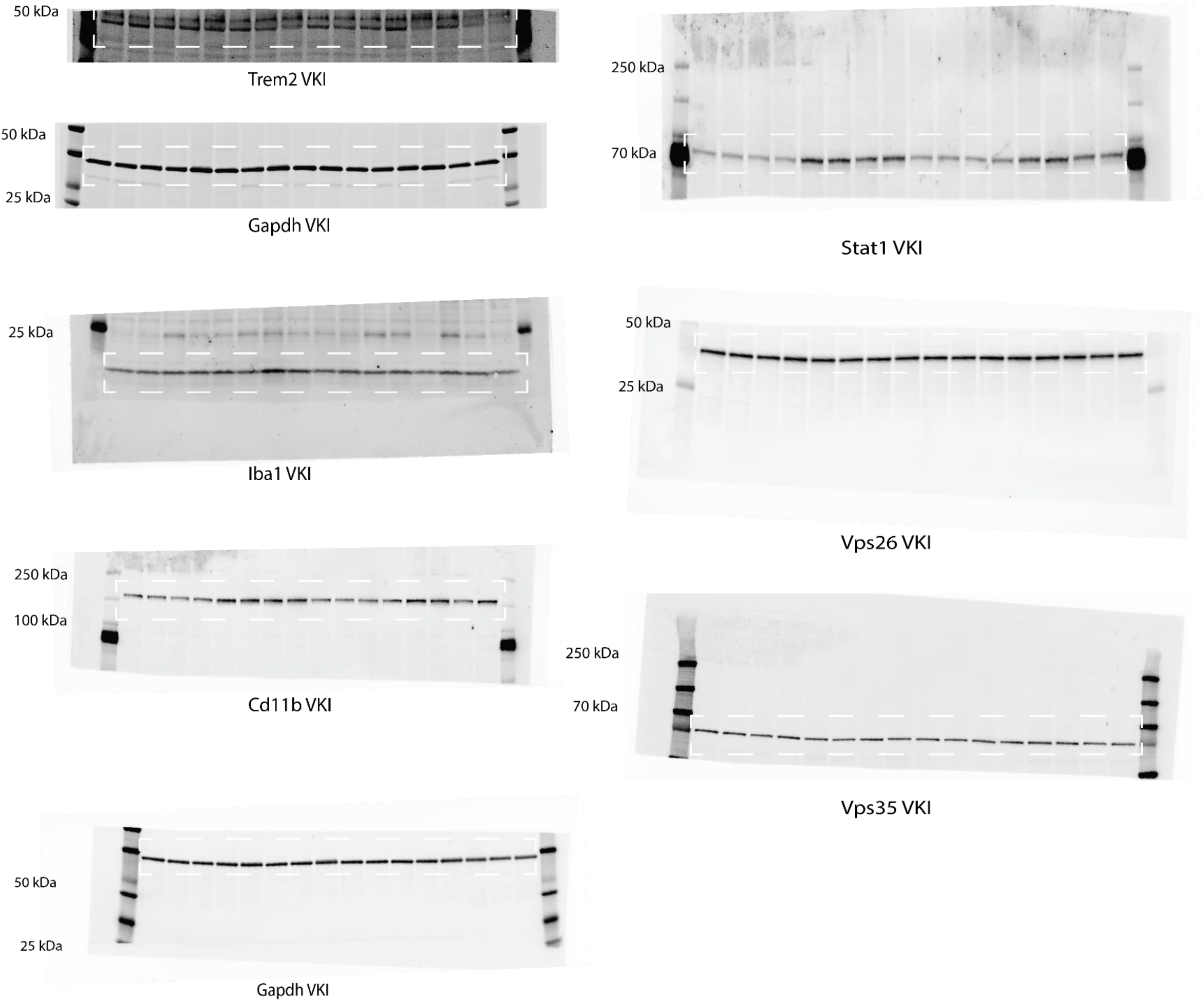

**Fig. S10.**
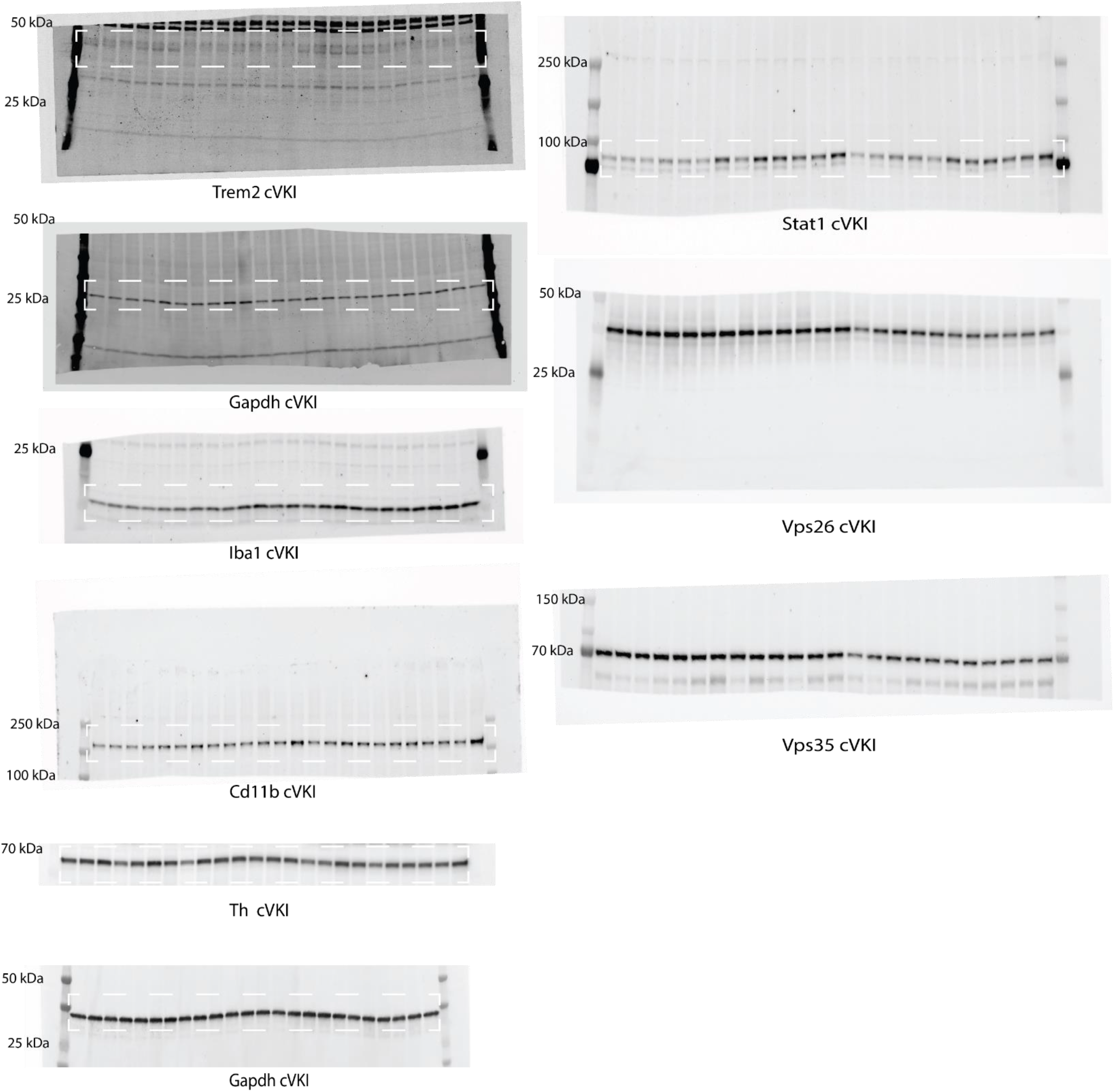

**Supplemental Table 1:** List of antibodies and dilutions.

| **Antibody** | **Source** | **Species** | **IHC** | **WB** |
| --- | --- | --- | --- | --- |
| Homer1 (VesL 1, Syn 47) | Synaptic Systems, 160 003 | Rabbit | 1:250 |  |
| lipocalin-2/NGAL | Proteintech, 31721-1-AP | Rabbit | 1:250 |  |
| Iba1 | Synaptic Systems, 234 308 | Guinea pig | 1:500 |  |
| bassoon | Synaptic Systems, 141004 | Guinea pig | 1:250 |  |
| Cd68 | AbD Serotec, MCA1957 | Rat | 1:500 |  |
| S100a9 [2B10] | Abcam, ab105472 | Rat | 1:250 |  |
| lipoprotein Lipase (LPL.A4) | Abcam, ab21356 | Mouse | 1:300 |  |
| Trem2 | R&D Systems, AF1729 | Sheep | 1:250 |  |
| Alexafluor IgG secondaries | Invitrogen |  | 1:500 |  |
| Iba1 | Wako, 016-20001 | Rabbit |  | 1:1000 |
| Trem2 | Cell signaling, 76765 | Rabbit |  | 1:500 |
| Gapdh | Thermo-Fisher Scientific, MA5-15738 | Mouse |  | 1:5000 |
| Stat1 | Cell Signaling, 9172S | Rabbit |  | 1:2000 |
| Th | Sigma, T2928 | Mouse |  | 1:1000 |
| Cd11b | Abcam, ab133357 | Rabbit |  | 1:2000 |
| Vps35 | H00055737- M02 | Mouse |  | 1:2000 |
| Vps26 | Abcam, ab23892 | Rabbit |  | 1:2000 |

